# Erythropoietin reshapes the adult hippocampal chromatin landscape to promote neurogenesis

**DOI:** 10.1101/2025.10.13.682070

**Authors:** Umut Cakir, Melanie Fritz, Umer Javed Butt, Riki Kawaguchi, Daniel Geschwind, Klaus-Armin Nave, Hannelore Ehrenreich, Manvendra Singh

**Affiliations:** Clinical Neuroscience, Max Planck Institute for Multidisciplinary Sciences, City Campus, Göttingen, Germany; Georg-August-University, Göttingen, Germany; Experimental Medicine, Department of Psychiatry and Psychotherapy, Central Institute of Mental Health, Medical Faculty Mannheim, Heidelberg University, J 5, Mannheim, Germany; Center for Neurobehavioral Genetics, Semel Institute for Neuroscience and Human Behavior, University of California Los Angeles, Los Angeles, CA, USA; Institute of Precision Health, University of California Los Angeles, Los Angeles, CA, USA; Department of Neurogenetics, Max Planck Institute for Multidisciplinary Sciences, City Campus, Göttingen, Germany; Department of Neuropathology, Charite, Berlin, Germany

**Keywords:** EPO, TE, pyramidal neurons, therapy, cognition, neuropsychiatric disorders

## Abstract

Understanding the molecular mechanisms by which erythropoietin (EPO) acts as neurotrophic factor that enhances hippocampal function and learning is essential to harness its therapeutic potential. Here, we employ single-nucleus ATAC-seq and RNA-seq to map the epigenomic and transcriptional landscapes of adult mouse hippocampus under recombinant human EPO (rhEPO) treatment. We discover significant lineage-specific chromatin remodeling predominantly in newly formed and immature excitatory neurons, highlighting a robust EPO-driven neurogenic response as the first direct evidence that an extrinsic factor can induce adult hippocampal neurogenesis. Notably, many EPO-induced accessible regions overlap ancient transposable elements, particularly ancient LINEs and SINEs, that are bound by key neurogenic transcription factors such as NEUROD1/2, NEUROG2, FOXG1, and ASCL1 and are linked to nearby genes governing neuronal differentiation and synaptic plasticity. Our findings uncover a previously unrecognized transposon-mediated mechanism underlying EPO-induced neurogenesis, highlighting an underappreciated role for TE-derived sequences in this process, and establish a publicly available single-nucleus multiomic atlas as a resource for understanding cell-type-specific gene regulation and neuroplasticity in the adult brain.

## INTRODUCTION

Erythropoietin (EPO) is a hypoxia-inducible cytokine best known for its indispensable role in erythropoiesis^1^. In recent years, however, it has become evident that EPO is expressed by multiple cell types in the adult brain, with particularly high levels in hippocampus, where it has been implicated as a pleiotropic neurotrophic factor that can modulate synaptic plasticity, neuronal survival and circuit remodelling^2^. Indeed, there is growing therapeutic interest in EPO for neuropsychiatric and neurodegenerative conditions^3^. Recombinant human EPO (rhEPO) improves cognitive performance in several patient groups and confers neuroprotection in preclinical models of neurodegeneration and brain injury^4^, supporting the existence of an endogenous “neuro-EPO” system in the adult central nervous system^2,5^. At the molecular level, several neuronal and glial lineages endogenously express EPO and its canonical receptor (EPOR)^6,7^, suggesting that EPO signalling is embedded in local cellular networks rather than restricted to the classical hematopoietic axis.

The hippocampus is a heterogeneous structure composed of excitatory and inhibitory neurons, progenitor populations, and diverse glial and vascular cells. Their contributions to learning, memory and mood are controlled by cell-type-specific gene regulatory programs. Bulk genomics has revealed that EPO treatment modulates plasticity-related gene expression and improves hippocampus-dependent behaviour, but lacks the resolution to connect these changes to defined cell types and cis-regulatory elements. Recent single-cell transcriptomic work demonstrated that rhEPO “re-wires” hippocampal networks, promoting differentiation and integration of pyramidal neurons while dampening inhibitory interneuron activity^8,9^. These studies, together with electrophysiological evidence showing enhanced excitatory and reduced inhibitory input onto newly integrated CA1 neurons, argue that EPO can reshape hippocampal circuitry. At the same time, EPO induces plasticity-associated genes and chromatin regulators, including Bdnf, Gap43, Psd95, Egr1 and Ep300, and alters HDAC5 localisation, pointing towards a major epigenomic component^3,8,10^. However, how EPO signalling impinges on the chromatin landscape of specific hippocampal cell types, and how this relates to adult neurogenesis, has not been resolved.

Single-nucleus assay for transposase-accessible chromatin (snATAC-seq) enables genome-wide profiling of chromatin accessibility at single-cell resolution and thus the identification of candidate cis-regulatory elements (cCREs): promoters and distal enhancers that underpin cell-type-specific transcriptional programs^11^. A key open question is how an extrinsic, clinically relevant modulator such as EPO reshapes chromatin accessibility in adult hippocampus *in vivo*, and which cell types and regulatory modules are most responsive. The snATAC-seq on hippocampi of mice treated with rhEPO allows us to capture dynamic regulatory events in rare hippocampal subpopulations and define the cCREs engaged by EPO, providing critical insights into how it reprograms neurogenic gene expression^8,12,13^.

An additional layer of complexity comes from the pervasive presence of transposable elements (TEs) in mammalian genomes. Roughly half of the mouse and human genome consists of TEs, including LINEs, SINEs, endogenous retroviruses (ERVs) and DNA transposons^14^. Far from being inert, TEs are increasingly recognised as to have been co-opted into gene regulatory networks in development, immunity and the brain^14–17^. In fact, a substantial subset of brain-specific cCREs is derived from TEs. For instance, in the human cortex, around 80% of recently gained (human-specific) regulatory elements overlap TEs^18^. Notably, species-specific ERVs, SINEs, and even the DNA transposons have been shown to serve as cCREs in neural progenitors, providing novel binding sites for neurogenic TFs^19,20^. Whether adult hippocampal neurogenesis induced by an extrinsic stimulus such as EPO also engages TE-derived regulatory sequences has not been addressed yet.

Here, we combine snATAC-seq with matched single-nucleus RNA-seq (snRNA-seq) and external TF ChIP-seq datasets to map the epigenomic and transcriptional consequences of sustained rhEPO treatment in the adult mouse hippocampus. We first construct an integrated single-cell atlas of chromatin accessibility and gene expression across major hippocampal lineages under placebo and rhEPO conditions. We then define EPO-responsive differentially accessible regions (DARs) globally and in a lineage-resolved manner, and relate these to changes in gene expression, TF motif usage and GRNs, with particular focus on newly formed versus mature pyramidal neurons. Finally, we interrogate the contribution of TE-derived sequences to EPO-responsive cCREs and ask whether they are preferentially engaged by neurogenic TFs.

Our data show that rhEPO induces extensive, cell-type-specific chromatin remodelling in adult hippocampus, with the most pronounced accessibility gains in newly formed excitatory neurons. These chromatin changes are coupled to transcriptional activation of genes involved in neuronal differentiation, synaptogenesis and synaptic plasticity, and are organised into regulons driven by canonical neurogenic TFs. A sizeable subset of EPO-responsive cCREs overlap TE insertions, particularly ancient LINE and SINE families, and these TE-derived sites are enriched for binding by neurogenic TFs in independent ChIP-seq datasets. Together, these findings place genome-wide chromatin accessibility changes rather than transcription alone at the centre of EPO’s pro-neurogenic action in adult hippocampus and suggest that TE-derived sequences form part of the regulatory substrate that can be engaged by the EPO-induced pro-neurogenic chromatin-regulation mechanism. Beyond the mechanistic insights, the snATAC-seq atlas presented here provides a resource for exploring stimulus-responsive regulatory landscapes in the adult brain.

## RESULTS

### Single-cell chromatin accessibility profiling uncovers 35 cell clusters in adult mouse hippocampus

To discern how rhEPO impacts chromatin accessibility in adult hippocampus, we performed snATAC-seq on the right hippocampus of 8 adult C57BL/6N mice (4 rhEPO-treated, 4 placebo-treated) following an established dosing paradigm (11 injections every other day starting from P28 to P49)^21,22^. Libraries were generated using the 10x Genomics Chromium platform and processed with Cell Ranger ATAC. After stringent quality control (fragment number, TSS enrichment, nucleosome signal, blacklist fraction), ∼35,000 high-quality nuclei were retained (Extended Data Fig. S1A-B, Table S1). Standard snATAC QC metrics were robust: we observed strong transcription-start-site (TSS) enrichment and characteristic nucleosomal banding, with <5% of fragments mapping to ENCODE blacklist regions (Extended Data Fig. S2A-B).

We constructed a unified peak set across samples using Signac framework^23^, applied TF-IDF normalisation and latent semantic indexing (LSI), and integrated samples with Harmony to correct batch effects^24^ (Fig. 1A). Dimensionality reduction and graph-based clustering resolved 35 distinct accessibility-defined clusters, visualised by UMAP (Fig. 1B). To assign cell identities, we integrated the snATAC-seq dataset with matched snRNA-seq profiles from the contralateral hippocampi of the same mice. Anchoring ATAC-derived gene activity scores to RNA expression via CCA yielded a joint embedding in which snATAC and snRNA nuclei overlapped well (Fig. 1C), and label transfer provided high-confidence cell-type annotations.

**Figure 1.**
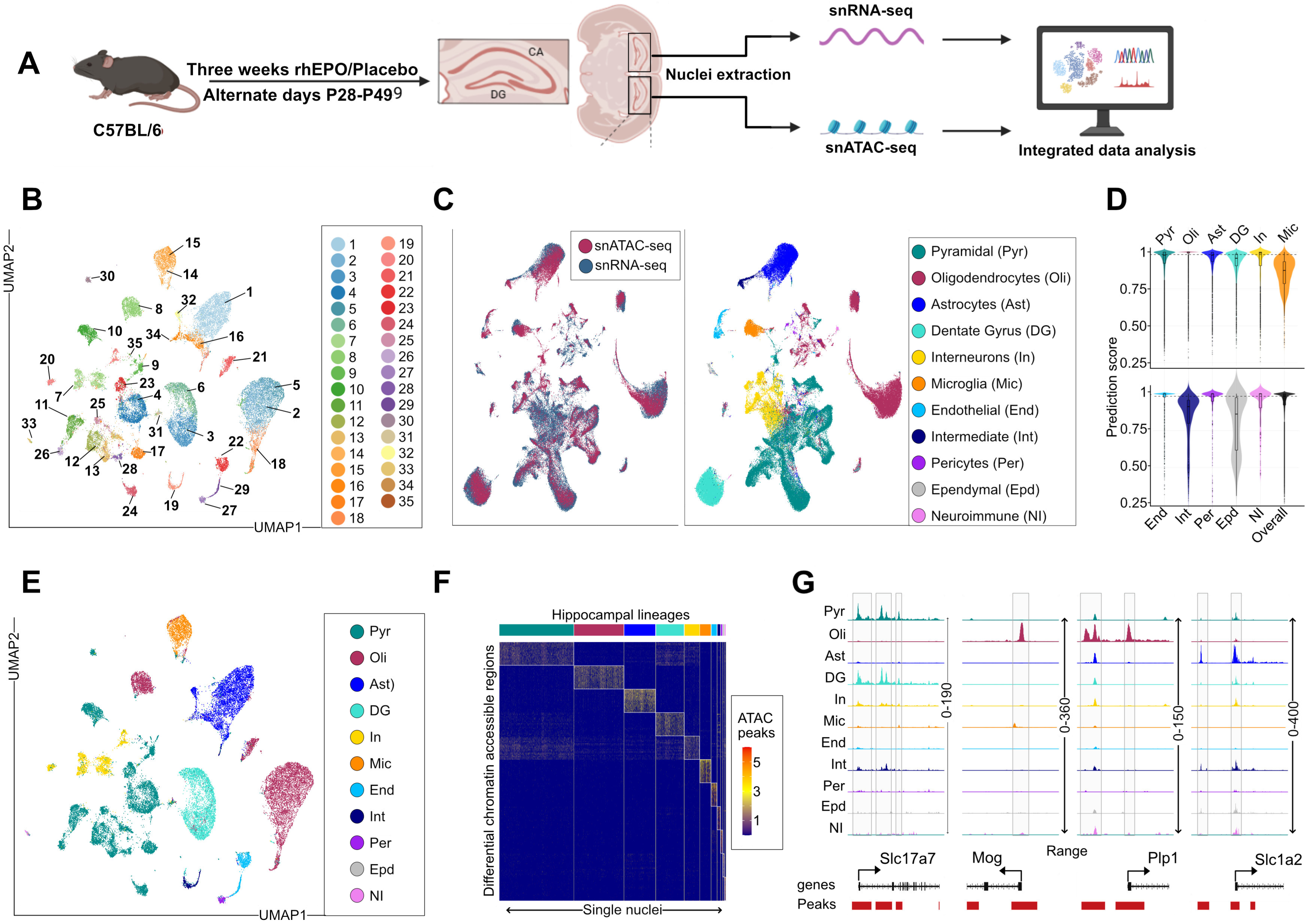
Integrated single-nucleus transcriptomic and chromatin-accessibility atlas of mouse hippocampus after rhEPO treatment. **(A)** C57BL/6 mice received recombinant human erythropoietin (rhEPO) or placebo every other day from postnatal day (P) 28 to P49. Hippocampal regions were dissected, and nuclei were isolated for single-nucleus RNA-seq (snRNA-seq) and single-nucleus ATAC-seq (snATAC-seq). The resulting datasets were jointly analysed to integrate gene expression and chromatin accessibility. Data processing included Cell Ranger peak calling, Signac-based quality control and preprocessing, Harmony batch correction, co-embedding snRNA-seq and snATAC-seq, cell-type annotation, differential accessibility testing, motif enrichment, gene regulatory network reconstruction, and identification of gene- and transposable element (TE)-associated candidate cis-regulatory elements (cCREs). **(B)** UMAP projection of the snATAC-seq dataset colored by 35 unbiased Seurat clusters. **(C)** Joint embedding of snATAC-seq and snRNA-seq using gene activity and expression. Left: UMAP projection of integrated nuclei, with snATAC-seq (red) and snRNA-seq (blue) showing strong overlap following CCA-based integration of ATAC-derived gene activity scores and RNA expression. Right: same embedding colored by major cell types assigned via label transfer and canonical markers, including pyramidal neurons (Pyr), dentate gyrus neurons (DG), interneurons (In), oligodendrocytes (Oli), astrocytes (Ast), microglia (Mic), endothelial cells (End), intermediate progenitors (Int), pericytes (Per), ependymal cells (Epd), and neuroimmune cells (NI). **(D)** Violin plots of prediction scores for cell-type assignments obtained by label transfer from snRNA-seq to snATAC-seq. Each violin represents a major hippocampal cell type, while the final column in the lower panel shows the overall distribution across all types. The dashed line marks the median overall prediction score, illustrating high cross-modality annotation accuracy. **(E)** UMAP projection of the snATAC-seq dataset, colored by cell-type identities transferred from the matched snRNA-seq reference shown in panel C. **(F)** Heatmap of differentially accessible chromatin regions (DARs, rows) across hippocampal cell types (columns). Scaled accessibility values (red/yellow = high, blue = low) highlight distinct cis-regulatory signatures for each lineage. For visualization, the top 20 peaks per cell type are shown. **(G)** Genome browser tracks showing chromatin accessibility at the Slc17a7, Mog, Plp1, and Slc1a2 loci across major hippocampal cell types. Highlighted regions mark cell-type-restricted accessible regions, illustrating lineage-specific regulatory landscapes.

We distinguished 11 major hippocampal lineages: dentate gyrus granule neurons, excitatory (pyramidal) neurons, interneurons, microglia, astrocytes, oligodendrocytes, endothelial cells, pericytes, intermediate progenitors, ependymal cells and neuroimmune cells (Fig. 1B-C). Accessibility at canonical marker loci supported these assignments: for example, astrocyte clusters displayed strong accessibility at Slc1a2, oligodendrocyte-lineage clusters at Plp1/Mbp/Mog or Pdgfra, dentate gyrus neurons at Prox1, and excitatory neurons at Slc17a7 (Fig. 1G, Table S2–S4). These chromatin-defined identities closely matched the transcriptomic clusters in the companion snRNA-seq dataset (Fig. 1C-E and Extended Data Fig. S3, Fisher’s Exact Test; p-value < 0.05), establishing a robust integrated atlas that we used as the basis for downstream analyses. Together, this integrated multi-omic atlas delineates the cell-type-specific regulatory landscape of the adult hippocampus, linking candidate cis-regulatory elements to gene expression programs. In line with this, distribution of peaks across hippocampal lineages further highlights distinct cis-regulatory signatures for each cell type (Fig. 1F). These integrated multi-omic analyses provide a foundation for constructing gene regulatory networks underlying hippocampal cell identity and the effects of rhEPO treatment. For instance, we observed that the *Slc1a2* (*EAAT2*) promoter is broadly accessible across multiple hippocampal lineages, whereas its distal enhancers are selectively open in astrocytes, yielding astrocyte-specific expression (Fig. 1G). This fits the established principle that promoters are often ubiquitously accessible while enhancer deployment is cell-type specific, as shown by ENCODE-scale promoter cCRE analyses^25^ and by astrocyte maturation studies documenting enhancer accessibility remodelling^26^.

### Characterization of hippocampal regulatory regions

To obtain an overview of regulatory diversity, we identified cell-type-specific DARs across the 11 lineages and performed motif enrichment analysis. We detected 135 TF motifs significantly enriched in accessibility peaks associated with specific clusters and their representative 11 lineages (log2 enrichment ratio > 2 and adjusted p-value < 1e^-10^), underscoring the breadth of regulatory programs operating in adult hippocampus (Fig. 2A and Extended Data Fig. S4, Table S2-3). As expected, well-known neurogenic and lineage-restricted TFs were prominent, including SOX TFs^27^, members of the EGR family^28^, the NEUROD1/2 and NEUROG1/2 factors, essential for neuronal differentiation^29,30^ and NRF1, a metabolic regulator that activates neuronal and synaptic genes^31^. Notably, other motifs highlighted TFs less emphasised in hippocampal biology but linked to neurodevelopment, such as GLIS1/2, ZBTB14, RFX2/5, the NR4A2::RXRA heterodimer, ZNF423, a zinc-finger TF^32^. These motif enrichments showed lineage-specific patterns: OLIG-like and SOX-family motifs were predominant in glial clusters, consistent with oligodendrocyte/astrocyte programs^33^, whereas EGR/ERG motifs dominated progenitor and early neuronal clusters except the interneurons, reflecting excitability-dependent genes and cell-cycle control in neural precursors^28,34^. In contrast, ETS/ELF motifs were highly enriched in endothelial and microglial clusters, aligning with their known association with immune regulatory programs^35^ (Fig. 2A and Extended Data Fig. S4, Table S2-3). This integrated atlas therefore captures both the cellular composition and the underlying regulatory codes of the adult hippocampus under control and EPO conditions.

**Figure 2.**
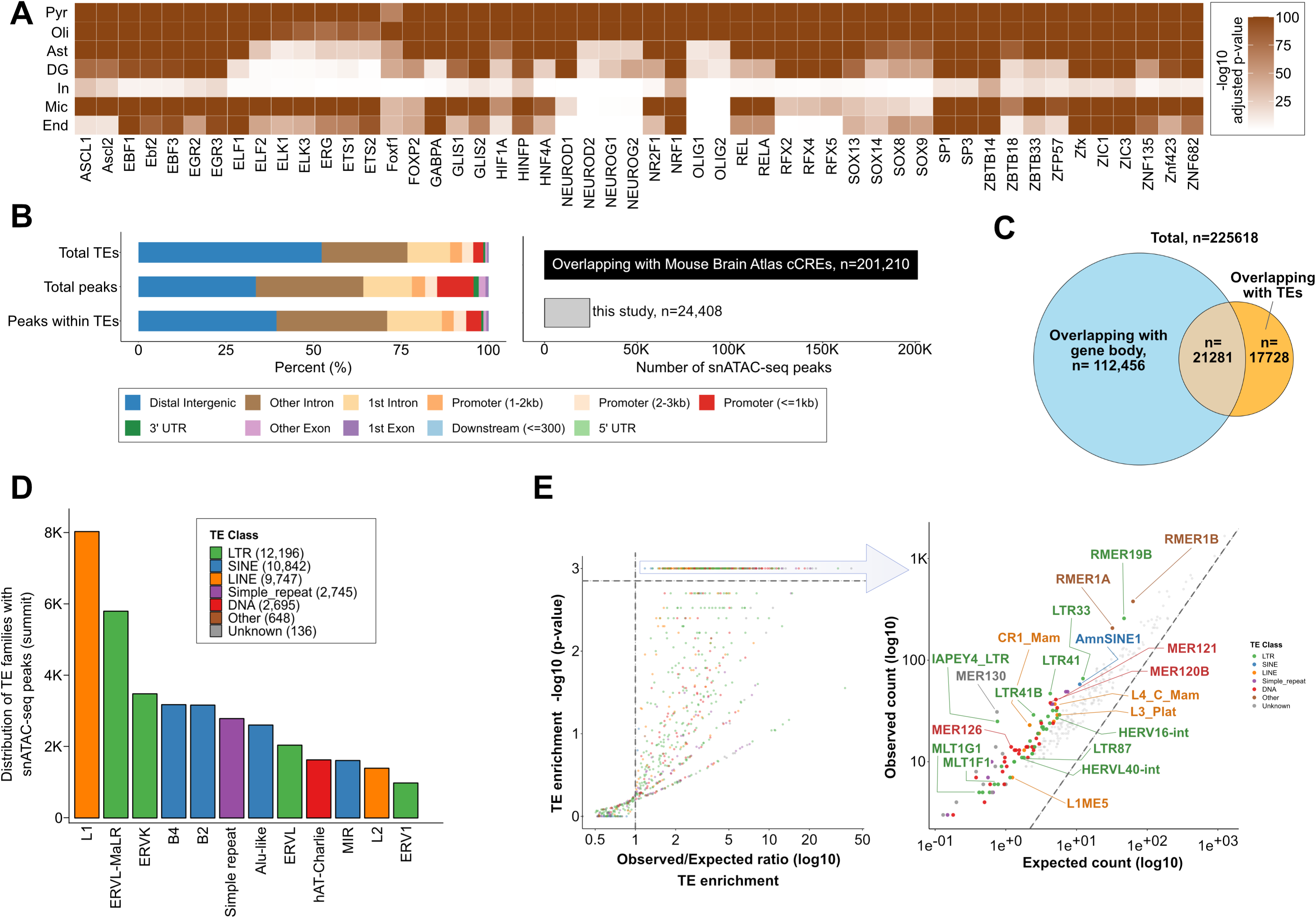
Cell-type TF motif usage and TE architecture of accessible chromatin. **(A)** Heatmap of significantly enriched transcription factor (TF) motifs (fold-enrichment >1.5; FDR <1×10⁻¹_J) in cell-type–specific DARs. Rows correspond to 7 major hippocampal cell types and columns to representative motifs (color scale = –log₁₀ adjusted p-value). The remaining 4 cell types are not shown due to limited motif detection resulting from their comparatively lower cell numbers. **(B)** Genomic distribution of (i) all annotated TEs in the mouse genome, (ii) all snATAC-seq peaks identified in this study (225,618 summits), and (iii) the subset of peaks overlapping TE sequences (39,009 summits). Bars indicate the percentage of features located in distal intergenic, intronic, exonic, promoter (<1 kb, 1–2 kb, or 2–3 kb), downstream, or UTR regions. Right: Overlap of snATAC-seq peaks with CATlas, showing 201,210 overlapping peaks (black) and 24,408 non-overlapping peaks (grey). **(C)** Venn diagram summarising the intersection between peaks that map to gene bodies (112,456), TEs (17,728), or both (21,281). **(D)** Bar chart of the TE families at peak summits, colored by TE class (LTR, SINE, LINE, simple repeat, DNA, other, and unknown). **(E)** The scatter plot shows log_2_(observed/expected) overlaps for each TE family versus –log₁₀ p-value. The right plot compares observed to expected counts for the top significantly enriched families highlighted in the left plot.

We next annotated snATAC-seq peaks by genomic context. Across all lineages, only ∼10.4% of accessible regions overlapped annotated promoters (±1 kb from TSSs), whereas 44.5% fell within introns, 36.9% in intergenic regions and 4.3% in exons or UTRs (Fig. 2B, Extended Data Fig. S5, and Table S4). This distribution is consistent with the notion that the majority of cCREs reside in intronic and intergenic space. Comparison with a recent single-cell atlas of adult mouse brain chromatin accessibility^36^ revealed that ∼10.8% of our hippocampal peaks were absent from prior annotations, indicating a set of unreported, potentially hippocampus-specific cCREs (Fig. 2B, Extended Data Fig. S5A, and Table S4). Given the prominence and the known contribution of TEs to regulatory landscapes, which constitute roughly half of the mammalian genome^17^ and are increasingly recognized as a major source of regulatory innovation^37,38^, we next intersected our snATAC-seq peaks with RepeatMasker annotations to quantify the extent of TE-derived regulatory activity. TEs frequently carry inducible enhancer or promoter activities, especially in the brain, where they have been shown to re-wire gene regulatory networks during development and evolution^19,37,39^. Approximately 20% of hippocampal cCREs overlapped annotated TEs, spanning LTR, LINE, SINE and other classes, consistent with previous reports that ∼25% of mammalian cCREs are TE-derived^39^ (Fig. 2C and Extended Data Fig. S5B). Notably, LTR elements exhibited disproportionately high accessibility, supporting the view that LTRs often function as self-contained regulatory modules for neighbor gene expression^37,39^ (Fig. 2D and Extended Data Fig. S5C). Among these, subfamilies such as ERVK, ERVL, RMER1A, RMER1B, IAPEY4, LTR41 and ORR1E showed the most significant enrichment in cCRE repertoires (Fig. 2D-E, Extended Data Fig. S5C, and Table S5). These TE-derived cCREs may serve as lineage-specific regulatory elements influencing hippocampal gene programs^40,41^. Together, these findings reveal that TEs are extensively integrated into the adult hippocampal regulatory network, providing a pool of fresh cell type enriched cCREs that likely contribute to hippocampal function.

### Epigenomic remodeling and enriched neuroplasticity regulators under rhEPO

To assess how rhEPO globally affects hippocampal chromatin accessibility, we compared aggregated peaks between EPO- and placebo-treated samples irrespective of cell identity. Logistic regression-based differential accessibility analysis identified 945 DARs (FDR < 0.01), of which 799 gained and 146 lost accessibility in rhEPO hippocampi (Fig. 3A). Upon mapping these DARs to annotated TSSs, we found that 639 of them were located within ±1 kb from TSS of known genes, out of which a substantial fraction of them overlapped with TE sequences (Fig. 3B). We then subjected these 639 genes to a gene ontology analysis, which indicated that these DAR-associated genes are significantly enriched in pathways for neuroplasticity, synapse organisation, and cognitive function, indicating a plausible mechanism of how EPO might re-wire the neurogenesis-associated transcriptional networks (Fig. 3C and Table S6). For instance, we detected robust EPO-induced chromatin accessibility at the Ascl1 promoter, a keytranscriptional regulator that promotes neural progenitor differentiation ^42,43^. Likewise, Egr2, an immediate-early gene of the EGR family is implicated in neuronal plasticity too^44^ (Fig. 3A). This finding is further supported by TF binding motif enrichment analysis of these DARs, which revealed a convergence on TF families that govern neurogenesis and synaptic plasticity (Fig. 3D). In addition to enhancer marking TFs SP1/2, rhEPO-open chromatin regions were highly enriched for binding motifs of Kruppel-like factors (KLFs), which is consistent with KLFs’ established role in neuronal maturation and associated with adult neurogenesis^45^ as well as early growth response factors (EGR1–3), which are immediate-early TFs critical for learning and memory^44,46,47^. Notably, motifs for AP-2 (TFAP2) factors, CREB/ATF family members (e.g. Atf1) and GLIS factors were over-represented (Table S5). These families have established roles in neuronal maturation, learning and memory ^48^, and activity-dependent gene expression^49^. Thus, at a global level, EPO promotes chromatin opening at promoter-proximal and distal sites that harbour motifs for neurogenic and plasticity-associated TFs.

**Figure 3.**
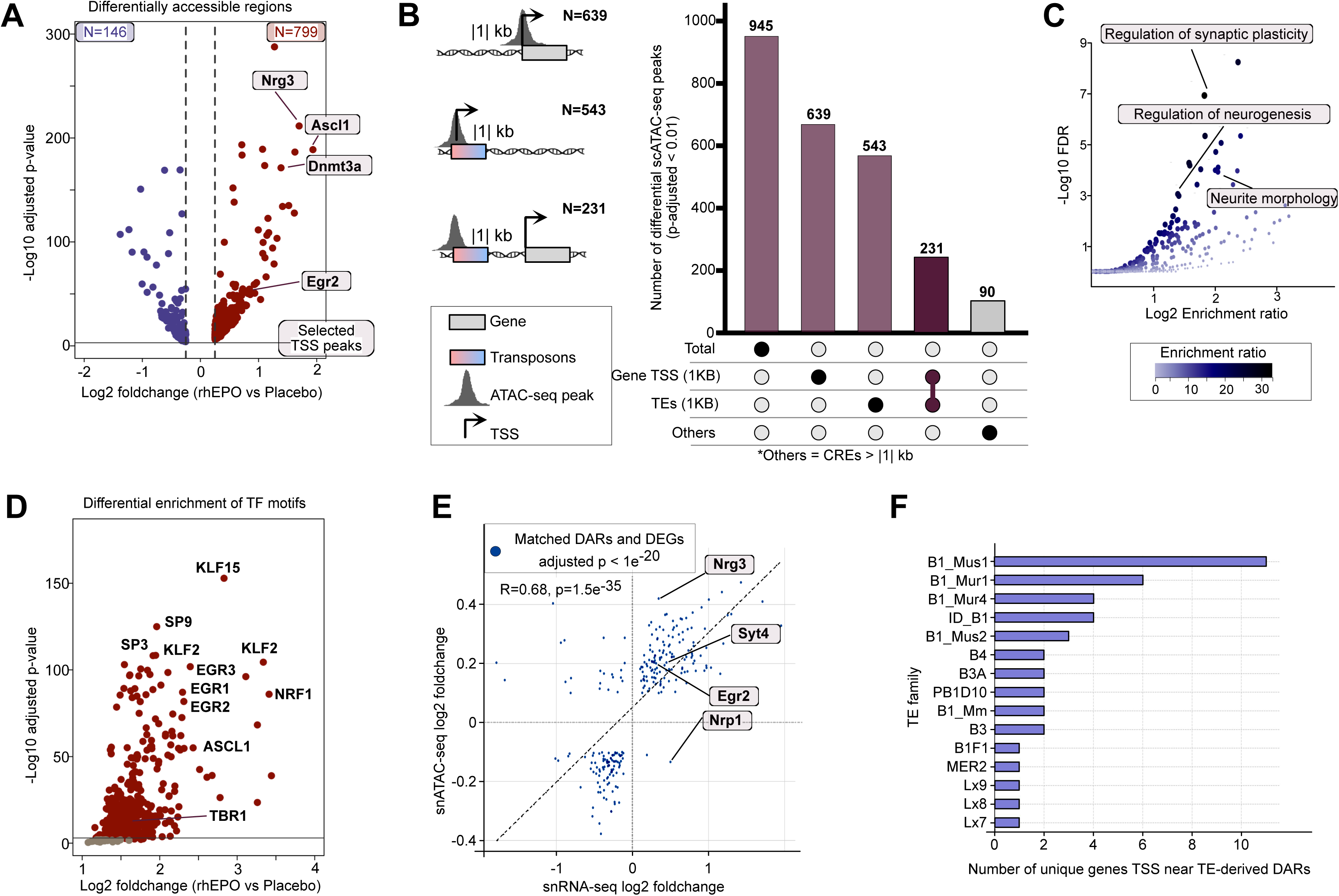
rhEPO remodels chromatin accessibility and engages neurogenic regulators. **(A)** Volcano plot of DARs between rhEPO- and placebo-treated hippocampi, identifying 945 DARs (799 gained, 146 lost). Neurogenic regulators (Ascl1, Dnmt3a, Nrg3, Egr2) are highlighted. **(B)** A total of 1,058 DARs (FDR < 0.01) were identified and grouped into 3 categories: promoter-proximal (within ±1 kb of a TSS), TE-associated (DARs overlapping TEs within ±1 kb of a TSS), and distal cis-regulatory elements (>500 bp from a TSS). The bar plot and accompanying UpSet diagram summarize the counts per category, with schematic illustrations on the left indicating classification criteria. **(C)** Gene ontology enrichment of DAR-associated genes, with significant terms related to synaptic plasticity, neurogenesis, and neurite morphology. **(D)** Motif enrichment analysis of DARs, revealing over-representation of TFs including KLFs, EGR1–3, ASCL1, NRF1, SP-family factors, and TBR1. **(E)** Scatter plot of differentially expressed genes from snATAC-seq inferred from gene activity and differentially expressed genes (DEGs, from snRNA-seq). Log_2_ fold changes (rhEPO vs placebo) show strong concordance (R = 0.68, p = 1.5 × 10⁻³_J), with representative neurogenic genes (Nrg3, Syt4, Egr2, Nrp1) highlighted. **(F)** Bar plot showing the number of unique genes with TSS located within ±1 kb of TE-derived DARs. Rodent-specific B1 SINE subfamilies (B1_Mus1, B1_Mur1, B1_Mur4, ID_B1) are the most frequent contributors, linking TE-derived accessibility to gene regulation.

Importantly, upon overlaying the similar analysis of differentially expressed genes (DEGs) from our snRNA-seq data, we observed a strong correlation between DARs identified by snATAC-seq and DEGs from our parallel snRNA-seq analysis, particularly for DARs proximal to DEGs (Fig. 3E). This correlation indicates that chromatin remodeling mediated by rhEPO precedes and is functionally concordant with transcriptional changes in hippocampal lineages. Our parallel snRNA-seq dataset too shows higher expression of Egr2, Syt4, Nrg3, Nrp1 and many other DAR-associated genes in EPO-treated hippocampus than in controls (Fig. 3E and Table S7)^8,9^. Several synaptic genes exhibited enhanced accessibility only in EPO-treated nuclei, such as Synaptotagmin-4 (Syt4), an activity-inducible vesicle protein that modulates BDNF release and learning-related synaptic plasticity, and Neuregulin-3 (Nrg3), a trophic factor that promotes excitatory synapse formation on interneurons^50,51^. We also found differential peaks at the Neuropilin-1 (Nrp1) locus, a receptor for axonal guidance cues (semaphorins). Thus, rhEPO application not only activates these pro-neurogenic and plasticity-related genes at the chromatin level, but also overexpresses their transcription, suggesting a coordinated epigenome-transcriptome response underlying EPO-induced alterations in the neural network.

Intriguingly, nearly half of the regions gaining accessibility with EPO treatment overlapped TE sequences, suggesting that EPO’s genome-wide epigenomic effects preferentially engage TE-derived loci. These TEs were predominantly rodent-specific SINEs of the B1/Alu and B2 families (including subtypes such as *ID_B1*, *B3*, *B1_Mus1/2*) (Fig. 3B and 3F), in line with emerging evidence that SINE-derived DNA and transcripts can serve normal neuronal functions^52^. More generally, SINE retrotransposons across the mammalian clade can recruit transcriptional regulators or RNA-binding proteins to nearby genes, thereby modulating gene expression in cis, as shown in primate, bovine and murine species^52–55^. The enrichment of B1/B2 SINE elements within EPO-responsive chromatin regions, located in close vicinity to DEG-associated loci (Fig. 3F), implies that the retrotransposon sequences may be tapped to mediate the regulation of gene programs activated by EPO.

### Lineage-Specific Chromatin Remodeling and TE-Driven GRN Activation in the Adult Mouse Hippocampus by EPO

To determine whether rhEPO exerts equal or lineage-specific effects on hippocampal chromatin states, we assessed differential chromatin accessibility across each of the 11 major hippocampal lineages provided in^8^ from the same set of mice. Using the logistic regression (LR) test with adjusted P < 0.01 threshold, we identified a total of 39,404 genome-wide DARs in all 11 lineages taken together. Although a subset of these DARs (n_J=_J 7,608) was common to multiple lineages, the majority (n_J=_J21,555) were lineage-specific (Fig. 4A and Table S8). While cataloguing the DARs in individual lineage, we found remarkably that pyramidal neurons alone accounted for approximately 50% (∼22,000) of the total DARs, highlighting their pronounced chromatin responsiveness to rhEPO treatment (Fig. 4B and Table S8). To assess the differential promoter activity, we tested what fraction of DARs in each of 11 lineages falls within 1-kb range of the TSS of annotated genes. We found that promoter-proximal DARs followed the same lineage-specific distribution pattern observed for DARs overlapping TE sequences across the hippocampal lineages. Interestingly, the distribution pattern of DARs among TEs and TSS between the 11 lineages was consistently proportional. Pyramidal neurons exhibited by far the greatest number of DARs on TE sequences (n = 3,897) and on the promoters of genes (n=1849), with an inclined bias toward increased chromatin accessibility, as 1065 TSS were upregulated versus 784 downregulated regions (Fig. 4B and Extended Data Fig. S6A, Table S8).

**Figure 4.**
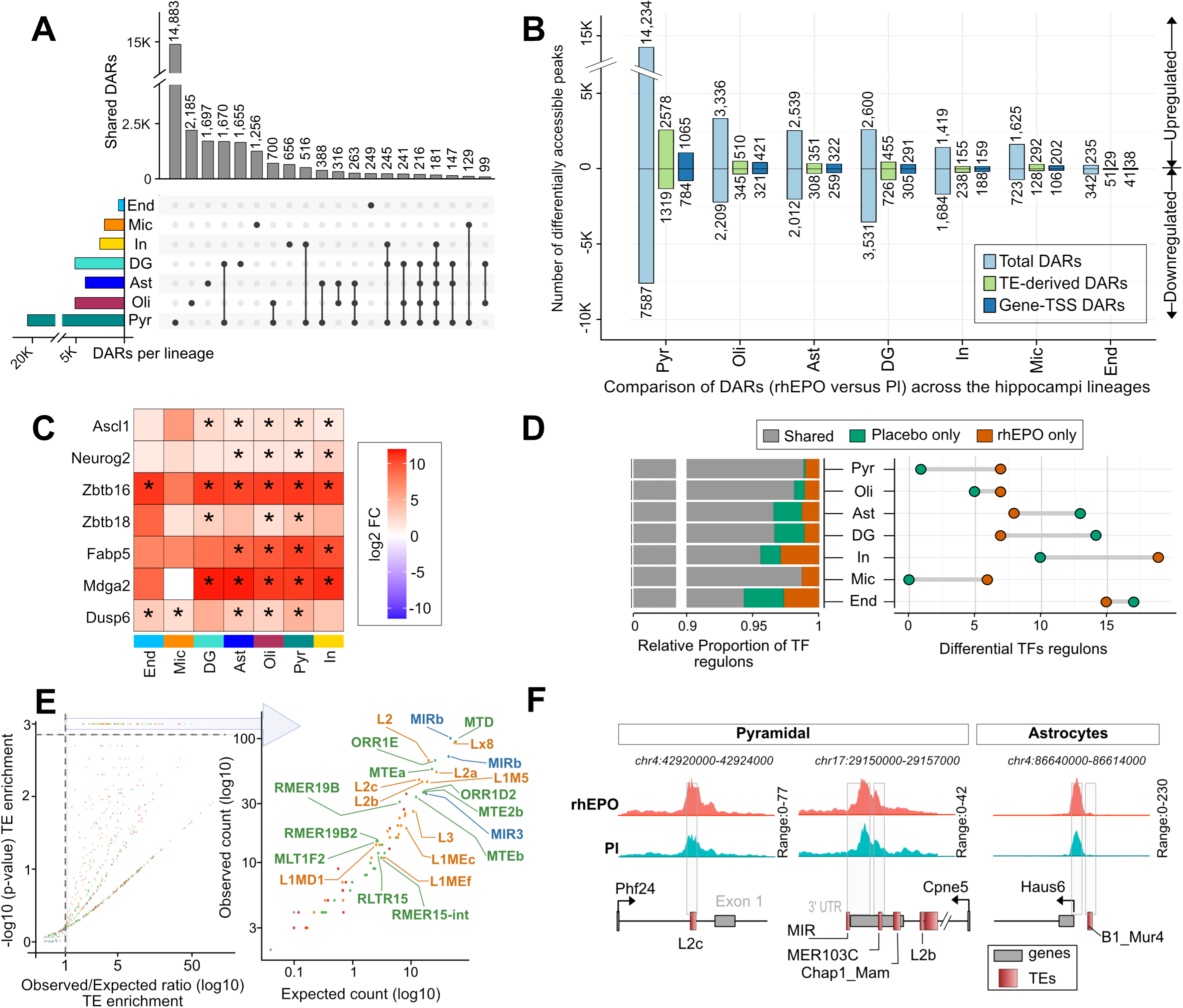
rhEPO selectively remodels chromatin accessibility, transposon usage, and TF programs across hippocampal cell types. **(A)** Global view of DARs (FDR < 0.01) between rhEPO- and placebo-treated nuclei. Horizontal bars indicate the number of DARs identified in each lineage (color legend, left); the UpSet matrix to the right shows how many DARs are unique or shared across lineages. **(B)** Stacked bars show, for each hippocampal lineage, the total number of DARs (light blue), the subset overlapping TEs (green), and the subset located within ±1 kb of gene TSSs (dark blue). Bars extending upward represent DARs with increased accessibility in rhEPO-treated samples, while downward bars indicate reduced accessibility compared with placebo. **(C)** Heatmap of rhEPO-induced promoter accessibility changes at selected neurogenic genes across hippocampal lineages. Colors indicate log_2_ fold change (EPO vs placebo), with red showing increased and blue decreased accessibility. Asterisks denote significant changes (FDR < 0.01). **(D)** Stacked bars show the proportion of regulons inferred by Pando that are shared, rhEPO-specific, or placebo-specific within each lineage. The dot plot on the right summarizes the absolute number of differential regulons, with orange indicating rhEPO-specific and green placebo-specific regulons. **(E)** TE-enrichment analysis for rhEPO-responsive DARs. The left scatter plot depicts log_2_(observed/expected) overlaps versus –log₁₀P for every annotated TE family (points colored by TE class; dashed boxes mark the top significantly enriched families). The right scatter plot compares observed to expected counts for the top significant families. **(F)** Genome browser snapshots of rhEPO-induced chromatin accessibility at TE-derived regulatory elements. In pyramidal neurons, increased accessibility was observed at an L2c element near Phf24 and at MIR and MER103C elements near Cpne5. In astrocytes, a B1_Mur4 element near Haus6 showed enhanced accessibility. Tracks display snATAC-seq signal for rhEPO (red) and placebo (blue), with genes (grey) and TEs (red) shown below.

Dentate gyrus showed the next highest DAR count (n = 6,131) but an opposite trend, as a majority of these regions had depleted accessibility for TEs (455 up vs. 726 down) and gene-TSSs (291 up vs. 305 down), whereas glial populations, oligodendrocytes (742 DARs), astrocytes (581 DARs) and Microglia (308 DARs) displayed moderate numbers of EPO-responsive TSSs, skewed slightly toward openness. In contrast, interneurons (347 DARs) exhibited the reverse pattern, with predominantly downregulated regions (159 up versus 188 down). Endothelial, Pericytes, neuroimmune, intermediate and ependymal cells were least affected (Fig. 4B and Extended Data Fig. S6A, Table S8). Thus, the majority of DARs in pyramidal neurons, microglia, and oligodendrocytes were upregulated, whereas dentate gyrus cells and interneurons showed predominantly downregulated DARs. Overall, this distribution pattern suggests that EPO’s chromatin remodelling effects are directed primarily toward excitatory lineage expansion and glial physiology, while concurrently limiting the accessibility changes associated with inhibitory neuronal programs.

We next tested a model, based on the above observations, in which EPO promotes open chromatin in excitatory neurons and glial cells by investigating if the key neurogenic gene promoter accessibilities are significantly upregulated in rhEPO samples. Expectedly, pyramidal neurons showed the activation of key neurogenic TFs and developmental regulators. For instance, Ascl1, Neurog2, and Pax6, the key factors driving neuronal commitment from precursors^56,57^, and Nrg3, a neuregulin promoting excitatory synaptogenesis^51^, all exhibited robust accessibility gains in pyramidal neurons, dentate gyrus, astrocytes, and oligodendrocytes (Fig. 4C and Extended Data Fig. S6B, Table S9). In addition to these established neurogenic genes, several less-characterised but pertinent regulators emerged among the top EPO-responsive promoters. For instance, the zinc-finger factors *Zbtb16* (Plzf) and *Zbtb18* (Rp58), both implicated in cortical neurogenesis and neuronal subtype development^58,59^, showed increased accessibility, as did *Fabp5*, *Mdga2*, and *Dusp6* (Fig. 4C and Extended Data Fig. S6B, and Table S9). The enrichment of these diverse gene promoters suggests that EPO activates a broad pro-neurogenic chromatin landscape, encompassing classical neurogenesis drivers and novel regulatory elements linked to neuronal maturation and synaptic integration.

Following these leads, we tested whether rhEPO treatment might differentially re-wire GRNs in hippocampal pyramidal and glial lineages. To systematically test this hypothesis, we applied the Pando algorithm, integrating our matched snRNA-seq and snATAC-seq datasets, to identify key TFs, their target regulatory regions, and downstream effector genes^60^. This analysis revealed that while most regulons were conserved between rhEPO and placebo conditions, approximately a dozen regulons displayed specific enrichment under rhEPO (Fig. 4D and Extended Data Fig. S7, Table S10). Expectedly, these rhEPO-enriched regulons were prominently driven by TFs involved in neuronal differentiation and maturation (Fig. 4D and Extended Data Fig. S7, Table S10). These observations support the model that rhEPO specifically engages neurogenic TFs to facilitate the elevation of gene expression essential for neuronal development and synaptic plasticity.

Additionally, we also found that a broad spectrum of TE families was significantly enriched within the DARs (Fig. 4E and Extended Data Fig. S8, Table S11). These enriched repeats included LTR retrotransposons of the ORR1E and MTD families, ancient LINEs L2/L2a and SINEs rodent B families B3, B4/B4A, the mammalian-wide interspersed repeat MIR/MIRb, which is an evolutionarily old SINE family^54,61–63^. This widespread TE activation is consistent with prior evidence that such elements can be co-opted as cCREs in neural cells^38,64,65^. Among many evidence of TE-derived cCREs, we observe increase accessibility of L2c, MER103C/MIR, and B1_Mur4 in rhEPO samples and seem plausible regulator of Phf24, Cpne5, and Haus6 genes, respectively (Fig. 4F). These patterns indicate that EPO treatment broadly unlocks repeat-rich genomic regions in a lineage-specific manner, suggesting that of TE families enriched in EPO-induced DARs act as cCREs that influence gene transcription in cis.

### Lineage-specific chromatin remodeling predominantly affects pyramidal neurons

We had shown that in rhEPO-treated mice, excitatory neuron lineage, particularly CA1 pyramidal cells were markedly over-represented, while the proportion of GABAergic interneurons was reduced^8,9^. Notably, we observed a dramatic expansion of newly formed pyramidal neuron subpopulations, with up to a 5-fold increase (∼500% of control levels) in rhEPO samples (Extended Data Fig. S9A-B). This dramatic expansion of immature excitatory neurons provides direct evidence that rhEPO treatment induces adult hippocampal neurogenesis, extending prior single-cell analyses showing ∼2-fold enrichment (∼200%) of immature pyramidal neurons and is consistent with earlier reports that prolonged EPO treatment increases mature CA1 pyramidal neuron numbers by ∼20%^8,13^. Even a single EPO dose is known to acutely elevate immature glutamatergic precursors within hours^13^. To further delineate how EPO orchestrates transcriptional networks underlying neurogenesis, we performed high-resolution chromatin accessibility clustering of pyramidal neuron lineages from our snATAC-seq data. Initial clustering from snRNA-seq identified 20 distinct pyramidal clusters^8^, whereas snATAC-seq revealed 29 distinct clusters (Extended Data Fig. S10A). Cross-modal integration allowed us to robustly annotate these clusters into 17 distinct cell types, showing substantial concordance between snRNA-seq and snATAC-seq modalities (Fig. 5A and Extended Data Fig. S10B). The two predominant pyramidal subgroups were newly formed (early-stage markers) and mature neurons (late-stage markers) that exhibited complete congruence between modalities, clearly segregating into discrete UMAP clusters. Following these leads, we next sought to characterize chromatin accessibility differences between newly formed and mature pyramidal neurons. Differential accessibility analysis identified 7,420 DARs distinguishing these two populations, of which 1,705 DARs mapped within gene promoters and 935 overlapped TE sequences (Fig. 5B and Table S12). Remarkably, ∼66 % of promoter-associated DARs displayed significantly higher chromatin accessibility in newly formed neurons, with gene ontology analyses revealing significant enrichment for neurogenesis and synapse formation (Fig. 5C). Of note, we observed that around 83% of DARs overlapping both promoter and TE-associated regions were upregulated in rhEPO samples.

**Figure 5.**
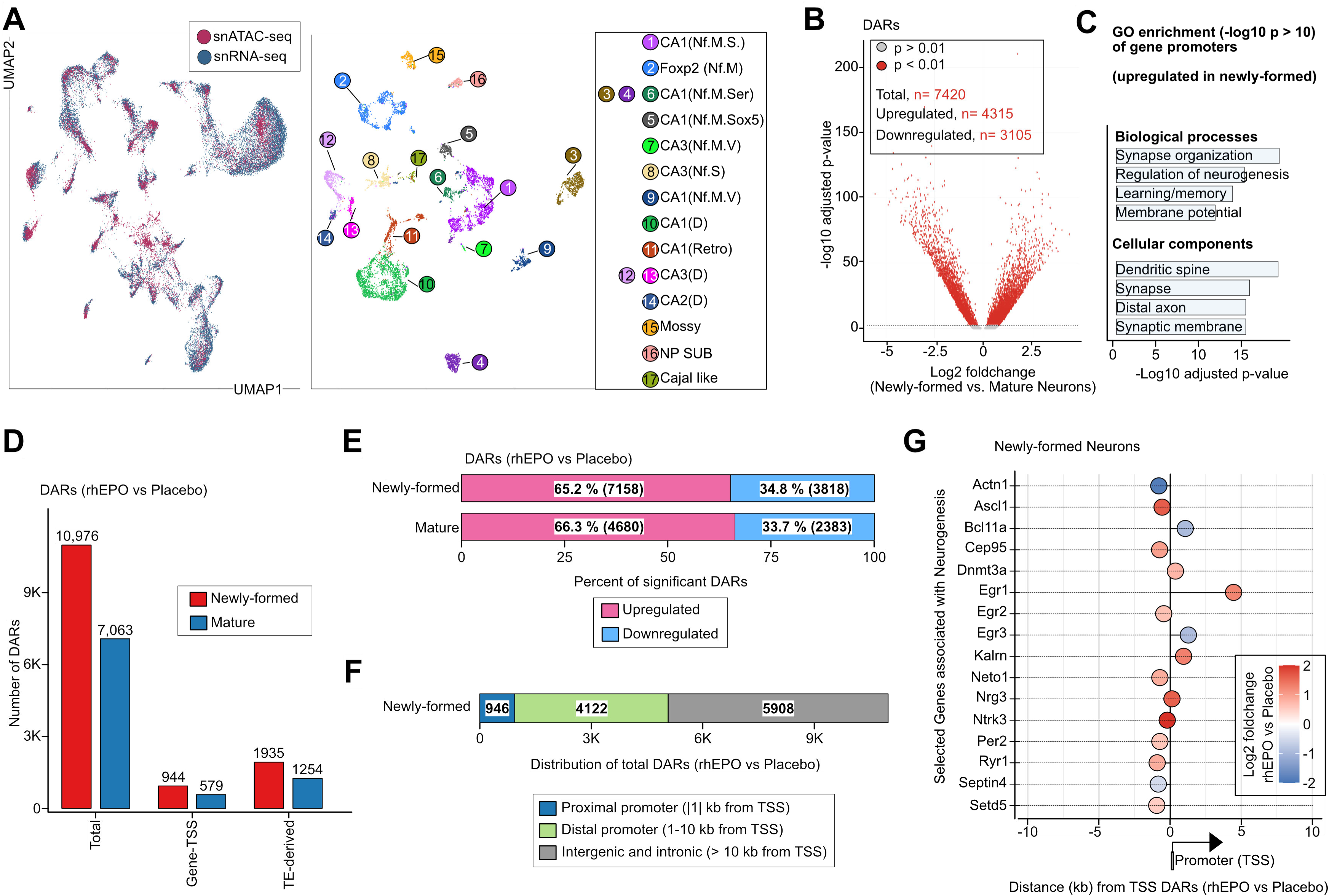
Pyramidal cell type-specific chromatin remodelling provoked by rhEPO. **(A)** Left: Joint UMAP embedding of pyramidal-lineage nuclei profiled by snATAC-seq (red) and snRNA-seq (blue). Right: UMAP of the same nuclei colored by clusters annotated through label transfer and marker gene activity, resolving newly formed neurons, mature pyramidal neurons, and other subtypes (e.g., mossy, NP SUB, Cajal-like cells). **(B)** Volcano plot of DARs between newly formed and mature pyramidal neurons (FDR < 0.01). A total of 7,420 DARs were identified, including 4,315 with increased accessibility and 3,105 with reduced accessibility in newly formed neurons. **(C)** Gene ontology enrichment analysis of promoter DARs upregulated in newly formed neurons. Significant terms (-log10p > 10) highlight processes linked to synapse organization, neurogenesis, learning, memory, and membrane potential and neuronal compartments (dendrite, synapse, distal axon, and synaptic membrane). **(D)** Numbers of DARs in newly formed and mature neurons (rhEPO vs placebo), shown for all DARs (left), promoter-proximal gene-TSS DARs (middle), and TE-derived DARs (right). Newly formed neurons exhibit higher counts across all categories. **(E)** Proportional distribution of DARs from panel D. In both newly formed and mature neurons, the majority of DARs are upregulated (pink), while a smaller fraction is downregulated (blue), indicating that rhEPO predominantly increases chromatin accessibility. **(F)** Genomic distribution of EPO-responsive DARs in newly formed neurons relative to the nearest TSS. DARs are classified as promoter-proximal (≤1 kb, blue), distal promoter (1–10 kb, green), or intergenic/intronic (>10 kb, grey). **(G)** Distance of significant DAR summits to the nearest TSS for selected neurogenic and synaptic genes in newly formed neurons. Each dot represents the nearest DAR to a gene TSS, colored by log_2_ fold change (red = increased, blue = decreased accessibility).

We then investigated chromatin accessibility differences between rhEPO and placebo conditions within immature and mature pyramidal neurons. As expected, the mature pyramidal neurons exhibited limited chromatin remodeling (∼7,000 DARs) compared to newly formed neurons (∼11,000 DARs) with ∼65% upregulated and ∼35% downregulated in both cases (Fig. 5E and Extended Data Fig. S11A). These results suggest the diminished epigenetic plasticity potential of rhEPO upon neuronal maturation. Within newly formed neurons, alongside a plenty of promoter-proximal DARs (n=946, ±1 kb from TSS), we identified an additional ∼4,000 distal DARs (within ±10 kb of gene TSS), marking putative enhancers and distal cCREs responsive to rhEPO (Fig. 5F, Extended Data Fig. S11B and Table S13). These distal regions prominently were located adjacent to immediate-early genes (Egr1, Egr2, Egr3), known mediators of neuronal activation and synaptic plasticity, as well as key neuronal specification genes (Ascl1, Bcl11a, Nrg3), and epigenetic regulators (Dnmt3a, Setd5) (Fig. 5G). Collectively, these data suggest that rhEPO preferentially promotes broad epigenomic remodeling and neurogenic transcriptional programs in newly formed neurons, while inducing limited, targeted adjustments in mature neurons.

### TE-Linked Neurogenic Enhancer Activity on EPO-Induced Chromatin Remodeling

We subsequently examined whether the DARs induced by rhEPO overlapping TEs participate specifically in GRNs activated in newly formed neurons. In newly formed (immature) pyramidal neurons, we identified a substantially greater number of EPO-induced TE-derived differentially accessible regions (TE-DARs) compared to mature pyramidal neurons, at both promoter-proximal and distal genomic locations. Interestingly, mature neurons showed merely 1,253, whereas newly formed neurons featured 1,934 TE-DARs, with 844 located within a range of ±10 kb near gene promoters (Fig. 6A, Extended Data Fig. S11C-D and Table S13). To further investigate the regulatory context of these sites, we intersected the TE-DARs with ChIP-seq profiles for 12 neurogenic transcription factors (TFs) (FOXG1, PAX6, ASCL1, NEUROG1/2, NEUROD1/2, SOX2, TBR1, EGR1, FOXO3, FOS, and JUN). Using established permutation tests^66^, we found hundreds of TE-DARs bound by one or more of these TFs - far more than expected by chance (Fig. 6B, Fig. 7A-C, Extended Data Fig. S12 and Table S14). Among TFs, NEUROD1 and FOXG1 exhibited the most extensive binding across TE families (295 and 256 TE families, respectively,pval < 0.001). A core set of 24 TE families, including ancient LINEs (L2, L3), LTR retrotransposons (LTR33, MER130, MLT1 series), and SINE elements (AmnSINE1, MamSINE1), was commonly enriched across multiple neurogenic TFs (Fig. 6C, Extended Data Fig. S13). Notably, many of these TE families have previously been exapted as gene regulatory enhancers in the brain. For instance, MER130 elements harbor consensus NEUROD/NEUROG-binding motifs and have been shown to function as enhancers during mouse neocortical development^67^. Likewise, AmnSINE1, an ancient SINE family, has well-documented enhancer activity crucial for mammalian brain development^68^. Together, our findings indicate that TE insertions from specific families have been specifically co-opted as part of the cis-regulatory repertoire of neurogenic TFs in mammalian brain.

**Figure 6.**
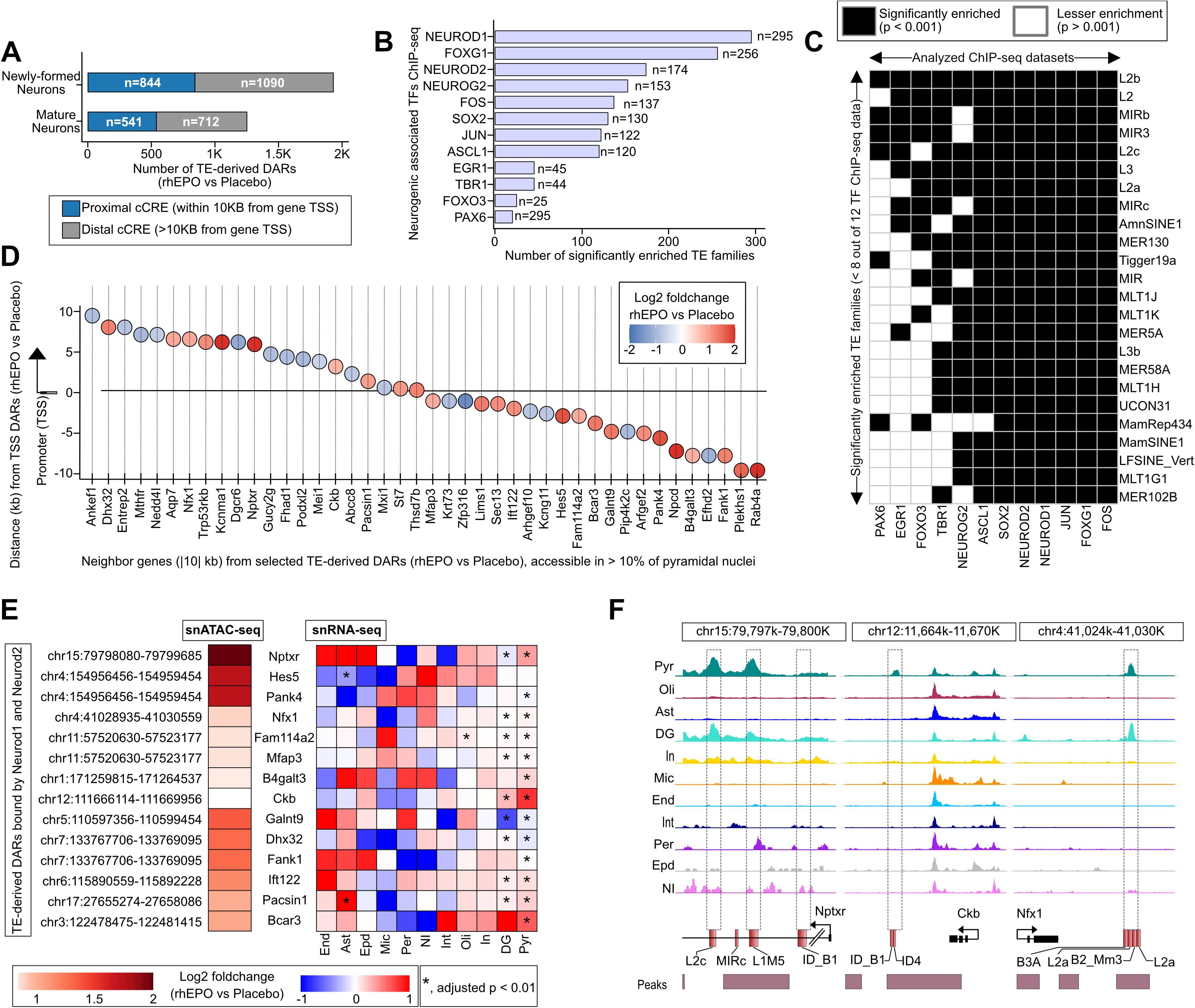
TE-derived DARs and their association with neurogenic TFs and gene regulation. **(A)** Numbers of TE-derived DARs (rhEPO vs placebo) in newly formed and mature neurons, separated into proximal cis-regulatory elements (cCREs within 10 kb of a gene TSS, blue) and distal cCREs (>10 kb from a gene TSS, grey). Newly formed neurons harbor more TE-derived DARs than mature neurons. **(B)** Bar chart ranking neurogenic TFs by the number of TE families significantly enriched within their ChIP-seq binding sites. Each bar represents the total count of enriched TE families associated with the indicated TF. **(C)** Matrix of significantly enriched TE families across TF ChIP-seq datasets. Black squares denote significant enrichment (p < 0.001), white squares denote no significant enrichment (p > 0.001). TE families such as L2b, L2, MIRb, and MIR3 are strongly associated with multiple neurogenic TFs. **(D)** Neighboring genes within ±10 kb of selected TE-derived DARs (rhEPO vs placebo) that were accessible in >10% of pyramidal nuclei. Each circle represents the closest DAR relative to the TSS, with position on the y-axis showing distance (kb) from the TSS, and color indicating log_2_ fold change (red = increased, blue = decreased accessibility under rhEPO). **(E)** Concordance between TE-derived DARs bound by neurogenic TFs and their nearest genes. Left, genomic coordinates of DARs identified by snATAC-seq, with color scale indicating log_2_ fold change (rhEPO vs placebo). Right, heatmaps of matched gene expression from snRNA-seq across hippocampal cell types. Asterisks denote significant transcriptional changes (FDR < 0.01), showing coordinated chromatin accessibility and expression responses to rhEPO. **(F)** Genome browser snapshots showing TE-derived DARs near neurogenic genes. Representative loci include Nptxr, Ckb, and Nfx1. Tracks display chromatin accessibility across major hippocampal lineages, with dashed boxes marking TE-overlapping elements that show lineage-restricted accessibility. Genes are shown in black, with TEs highlighted in red.

**Figure 7.**
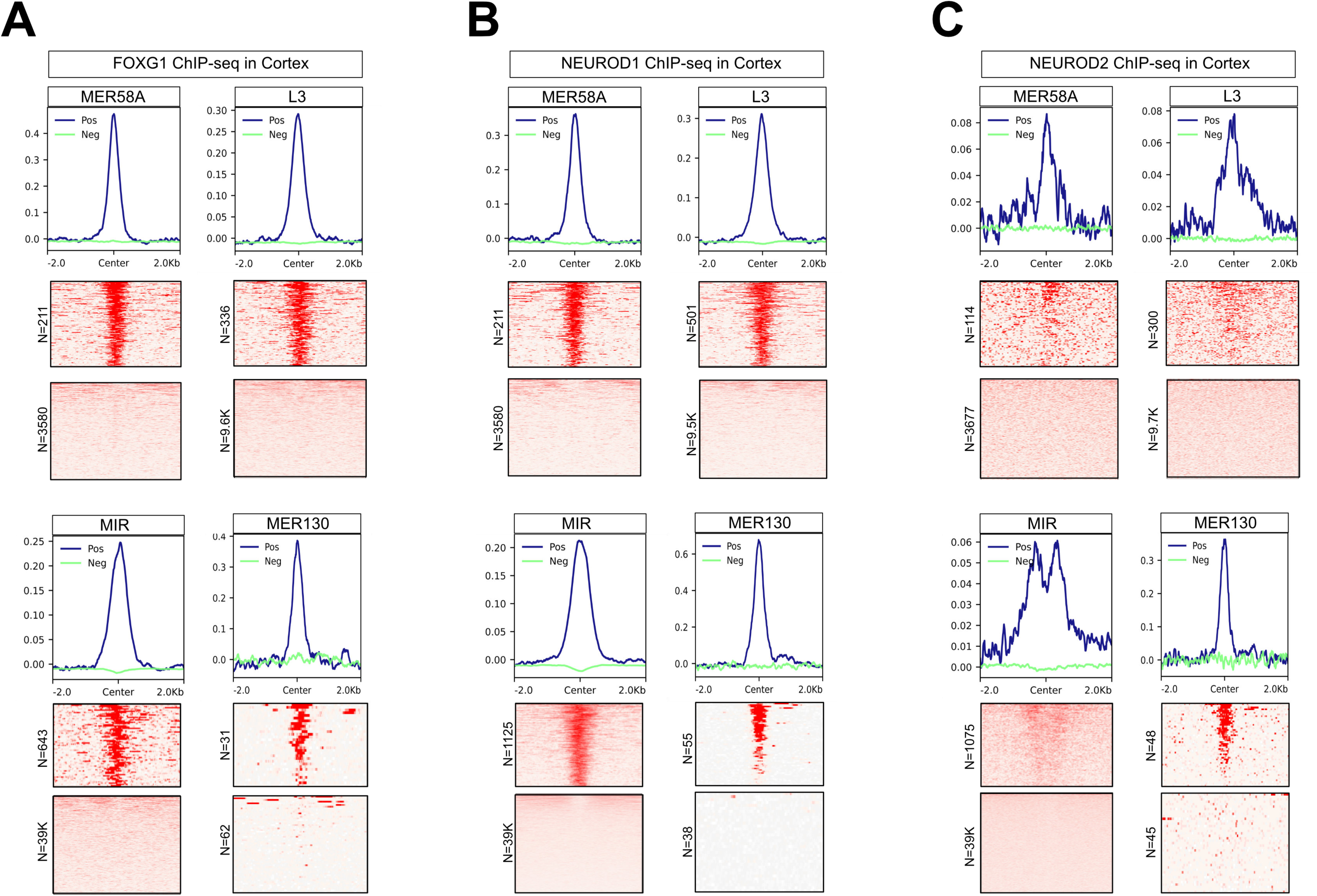
Neurogenic transcription factors FOXG1, NEUROD1, and NEUROD2 exhibit selective binding to transposable-element–derived sequences. **(A)** FOXG1 ChIP-seq shows aggregate ChIP-seq signal profiles (top) and corresponding heatmaps (bottom) centered on representative repeat element families (MER58A, L3, MIR, MER130). “Positive” (Pos, blue) represents transposable elements overlapping FOXG1 ChIP-seq peaks, whereas “Negative” (Neg, green) indicates transposable elements without peaks. Each heatmap displays signal density within ±2 kb of the element center, revealing distinct FOXG1 enrichment over specific TE subfamilies. **(B)** NEUROD1 ChIP-seq shows aggregate signal profiles and heatmaps demonstrating pronounced NEUROD1 binding across the same TE families (MER58A, L3, MIR, MER130). **(C)** NEUROD2 ChIP-seq shows meta-profiles and heatmaps depicting NEUROD2 binding over the same TE families. Altogether, these analyses demonstrate that ancient transposable-element subfamilies function as conserved cis-regulatory modules co-opted into the neurogenic transcriptional program.

To explore the functional relevance of these TE-derived DARs bound by neurogenic TFs, we examined the genes located nearby. We found 42 TE-containing DARs in rhEPO versus placebo comparison (filtered by >10% cells accessible at least in one condition) within 10 kb of range from genes TSS (Fig. 6D and Table S15). Consistent with our thesis, gene ontology analysis showed that the nearest genes to TF-bound TE-DARs were highly enriched for biological processes related to neurogenesis and synaptic integration (Extended Data Fig. S13 and Table S16). The genes nearest to these TE-DARs have well-documented roles in neuronal differentiation and synapse development, such as Hes5, Kcnma1, Pacsin1, Ankef1, Rab4a, Nptxr, and Nedd4l (Fig. 6D-E). These results demonstrate that ancient TE sequences serve as regulatory elements specifically during the developmental window of new neuron formation, thus enhancing the expression of genes critical for neuronal integration and synaptic plasticity. Furthermore, chromatin accessibility at these TF-bound TE-DARs was markedly increased in rhEPO samples (FDR < 0.01), which was accompanied by concordant transcriptional upregulation of nearby genes, specifically in newly formed neurons and dentate gyrus (Fig. 6E). For instance, our selected aforementioned candidates were exclusively accessible in rhEPO newly formed neurons, potentially driving the expression of several genes, as these genes are upregulated in our snRNA-seq data (Fig. 6E-F). Together, leveraging publicly available ChIP-seq datasets, we confirmed that canonical neurogenic TFs extensively bind the EPO-induced TE-derived enhancers *in vivo*. Crucially, many of these TF-occupied transposon insertions lie adjacent to genes involved in neuronal differentiation and synapse formation, which show concordant increases in accessibility and expression under EPO treatment. In summary, our findings underscore an innovative regulatory mechanism whereby rhEPO engages TE-derived enhancers targeted by neurogenic TFs, facilitating epigenomic reprogramming essential for hippocampal neurogenesis and synaptic plasticity.

## DISCUSSION

In this study, we constructed a high-resolution atlas of chromatin accessibility in the adult mouse hippocampus and demonstrated that EPO, a traditionally hematopoietic hormone, can profoundly reprogram the adult neural epigenome to drive neurogenesis. This expands upon prior single-cell brain atlases^11,69^ by focusing on the hippocampus and incorporating an experimental perturbation (EPO), thereby providing new insights into how adult neurogenic niches can be modulated. Crucially, we found that EPO’s effects are highly cell-type-specific, predominantly impacting excitatory neuron lineages (pyramidal neurons), which showed the most pronounced chromatin remodeling and population expansion in response to the treatment.

### EPO as a potent extrinsic modulator of adult hippocampal neurogenesis

These findings enrich our understanding of adult hippocampal neurogenesis, a process first recognized in rodents in the 1960s^70,71^ and later confirmed in humans using classical and current state-of-the-art methods^72,73^. The adult hippocampus retains progenitor cells that can differentiate into new neurons mainly in response to intrinsic or extrinsic stimuli. While we lack the temporal data, our results provide the first direct demonstration that EPO serves as an extrinsic trigger of adult hippocampal neurogenesis, which sets a mechanistic layer to earlier observations that EPO improves cognitive function and enhances neuroplasticity^2,13,74^. *In vivo* studies in healthy mice and clinical trials in patients have shown that EPO can improve learning and memory independent of its role in erythropoiesis^2,75–77^. We recently reported that EPO treatment increases mature pyramidal neuron numbers by ∼20% and newly formed neurons by ∼200% without altering cell proliferation, suggesting EPO promotes differentiation or maturation of pre-existing neural precursors. In this study, we observed a substantially greater relative expansion (∼5-fold) of the young pyramidal neuron population under EPO. This striking difference may either reflect the sensitivity of our chromatin-based approach in capturing nascent neurons, as well as a possibly more robust EPO regimen, or a region bias in some samples. These hypotheses remain to be tested. Importantly, our data support the model that EPO drives the differentiation of progenitors rather than inducing widespread cell division, as we detected increased accessibility at neurogenic transcription factor loci (e.g., *Ascl1*, *Neurog2*, *Nrg3* etc.) but did not observe evidence of aberrant cell-cycle activation in non-neuronal lineages. This aligns with the notion that EPO biases cell fate choice toward neurons in the existing precursor pool. The net effect is an enrichment of pyramidal neurons at the cost of interneurons. Pyramidal neurons integrate in the hippocampal circuitry, leading to potential implications for mood, memory, cognitive enhancement and recovery from brain injury.

### TEs as a regulatory innovation in hippocampal gene regulation

Despite TE’s known roles in other brain regions, the contribution of TEs to regulatory landscapes in the adult hippocampus has remained unexplored at single-cell resolution. One of the major highlights from our study is the extent to which TEs contribute to the gene regulatory changes underlying the EPO-induced phenomenon in hippocampi. We found that roughly one-fifth of all open chromatin regions in the adult hippocampus are derived from TEs, even under basal conditions. This observation is in line with large-scale epigenomic surveys showing that a substantial fraction of brain-specific cis-regulatory elements are TE-derived^36^. Notably, the hippocampus, particularly the dentate gyrus, a hub of adult neurogenesis, has been reported to exhibit high levels of TE activity and mosaicism^73^. TEs are thought to create genomic variability in neural progenitors and even generate long non-coding RNAs in the brain that fine-tune gene expression^64,78^. Our results extend these findings by pinpointing specific TE families that function as accessible chromatin sites in distinct hippocampal cell types. For example, we observed that certain LINE, SINE, and endogenous retrovirus (ERV) families, such as RMER1 retrotransposons and IAPEY4/ORR1 endogenous retroviral elements, are enriched in open chromatin unique to hippocampal neurons. These TE-derived sequences appear to serve as cell-type-specific enhancers or promoters, an interpretation supported by their overlap with known neuronal gene loci and motif enrichments.

TEs are increasingly recognized as drivers of gene regulatory network evolution. Many TEs carry binding site motifs for transcription factors and can thereby introduce novel regulatory modules. Classic examples include the co-option of a SINE element (*AmnSINE1*) as an enhancer essential for mammalian brain development^52,53^ and the MER130 DNA transposon family, providing binding sites for neural transcription factors in the developing cortex^64,65^. Our data demonstrate that similar TE-mediated regulatory mechanisms are active in the adult hippocampus. We found motifs for neural fate determinants e.g., NeuroD, Sox, and Klf, embedded within several TE sequences that are accessible in neurogenic cell clusters. EPO’s influence on the epigenome appears to leverage this TE repertoire. Remarkably, by aggregating all hippocampus cells, we observed that about 50% of the regions that gained DARs with EPO treatment were endogenous retroelement-derived sequences (Fig. 3), which is a proportion far exceeding their baseline representation. This enrichment suggests that EPO may activate a specific subclass of dormant regulatory elements to drive the nearby genes. One intriguing possibility is that these TE-DARs carry latent binding motifs for inducible transcription factors and become functional enhancers only under certain stimuli, as in this case, EPO stimulation. Such a mechanism would represent a previously unappreciated mode of gene regulation: the *de novo* recruitment of ancient genomic elements to drive a contemporary cellular response.

### Mechanisms of EPO-mediated neuroplasticity

Our integrative multi-omic analysis sheds light on the gene regulatory networks fine-tuned by EPO and how they intersect with TEs. By combining chromatin accessibility data with matched single-cell transcriptomes and leveraging a network inference algorithm, we identified key EPO-responsive regulons, which are sets of transcription factors and target genes that act in cascades. Prominently featured in these regulons were classical neurogenic transcription factors like NEUROD1, NEUROD2, FOXG1, PAX6, and JUN, known to orchestrate the progression of neural stem cells into mature neurons^79^. In EPO-treated hippocampi, these factors were not only upregulated at the mRNA level in pyramidal lineages, but their binding motifs were also significantly enriched in the EPO-gained accessible chromatin regions, and many of those were derived from TEs. This implies that EPO induces a feed-forward loop: it elevates neurogenic TF expression, and those TFs in turn bind to and activate TE-derived cCREs associated with neuron-specific genes, reinforcing the neurogenic program. Our incorporation of external ChIP-seq data from developing brain or neural culture systems provided orthogonal validation for this model, showing, for example, that NEUROD1/2 and NEUROG2 binding sites overlap many of the LINE/SINE/ERV-derived cCREs that EPO opened *in vivo*. While these findings resonate with reports that cell-type-specific TFs often exploit transposon-derived sequences for binding^80^, here we demonstrate this phenomenon in an active, inducible context within the adult brain.

Taken together, we propose a conserved evolutionary mechanism at play: ancient TEs embedded in the genome have been repurposed as cCREs in neural cells, and during an EPO-induced neurogenic stimulus, the brain taps into this reservoir of regulatory elements to rapidly reshape gene expression. In evolutionary terms, this represents an efficient strategy – rather than evolving entirely new gene regulatory sequences from scratch, genome has preserved a large bank of TE-derived cCREs that lie dormant until the appropriate transcriptional signals such as those triggered by EPO/EPOR signaling, activate them. The outcome is a robust transcriptional activation of neurogenesis-promoting genes. This concept aligns with the hypothesis that TEs facilitate the acquisition of novel cis-regulatory elements over evolutionary time^37,81^, and our data illustrate that such TE-facilitated regulatory innovation can be deployed within a single organism’s lifespan, enabling context-specific gene regulation to adapt to physiological stimuli.

### Implications of EPO-mediated re-wiring of GRNs in regenerative therapy

Our study carries broad implications. First, it highlights a previously underappreciated molecular pathway by which extrinsic factors like rhEPO can induce neuroplastic changes: through large-scale epigenomic remodeling of chromatin accessibility. This complements classical signaling and gene expression analyses by demonstrating that changes in the “accessibility” of the genome are a critical intermediate step linking external signals to sustained cellular phenotypes. Of note, EPO is known to be upregulated in the brain under hypoxic or injury conditions, and it has been considered a natural neuroprotective agent^77,82^. Our findings raise the possibility that part of the brain’s injury response may involve EPO-driven engagement of transposon-based cCREs to activate genes for repair, regeneration, or circuit reorganization. Second, our work highlights the functional importance of the vast repetitive genome in normal physiology. Whereas TEs have traditionally been studied in the context of genome instability or evolution, here we show they play a central role in acute, beneficial plasticity of the adult brain when challenged by external stimuli. This bridges the gap between transposon biology and neurobiology, suggesting that they are intimately involved in processes like learning, memory, and regeneration. Finally, from a translational perspective, identifying TE-derived cCREs that are activated by pro-neurogenic stimuli could open new therapeutic strategies. If certain transposon-derived cCREs are master switches for neurogenic gene programs, they might be targeted by gene editing or epigenetic drugs to stimulate neuron replacement in ageing or neurological disorders. Moreover, our single-cell multi-omic atlas of the EPO-treated and control hippocampus is publicly available as a resource for the community. It provides a catalogue of candidate regulatory elements for each major hippocampal cell type. In summary, our work not only provides the first mechanistic demonstration of EPO-induced adult hippocampal neurogenesis but also lays a foundation for future explorations into the “regulatory dark matter” of the genome in adult brain function and therapy.

## CONCLUSIONS

While our present study on rhEPO-mediated epigenomic reorganization in the adult hippocampus is profound, it also confers obvious limitations. Firstly, our analysis captures a snapshot of a single time point after rhEPO treatment, so we could not resolve the temporal dynamics of chromatin and transcriptional changes. While we identified a strong association between TFs-TEs interplay and chromatin accessibility as a proxy for regulatory activity maneuvered by EPO, the targeted perturbation e.g. CRISPR deletion of a candidate TE enhancer, and other epigenetic layers, such as histone modifications or 3D chromatin architecture, will be necessary to confirm the causality. Despite these limitations, our work reveals a compelling new paradigm in which an exogenous factor (rhEPO) leverages the genomic potential of transposable elements to execute a complex biological program such as neuronal development and plasticity. This insight expands our understanding of gene regulation in the adult brain and could inform novel strategies to enhance brain repair, cognitive function, and resilience against neurodegenerative conditions. By uncovering the latent regulatory power of transposons in neural plasticity, we pave the way for future studies to explore the therapeutic modulation of the “repeatome” in the service of regenerative medicine.

## METHODS

### Ethical approval

All mouse experiments were approved by the local Animal Care and Use Committee (Niedersächsisches Landesamt für Verbraucherschutz und Lebensmittelsicherheit, LAVES – AZ 33.19-42502-04-17/2393) and conducted in strict accordance with the German Animal Protection Law. Every effort was made to minimize the number of mice used and their suffering.

### Experimental model and treatments

Male C57BL/6N mice received 11 intraperitoneal injections of recombinant human erythropoietin (rhEPO, 5000 IU/kg body weight) or placebo (0.01 mL/g of solvent) every other day for 3 weeks, starting at postnatal day 28 (P28). In total, 23 male mice were used (rhEPO = 11, placebo = 12). Twenty-four hours after the final injection (P49), all animals were sacrificed by cervical dislocation. For each sample, 2 right hippocampi from mice of the same treatment group were pooled into a single tube (except for one rhEPO sample, which included only one hippocampus), resulting in 6 biological replicates per condition (N=6 rhEPO, N=6 placebo) for snRNA-seq (GSE220522). Likewise, the left hippocampi from 4 tubes of each group (N=4 rhEPO, N=4 placebo; tube labels are: A1, A2, A4, A6, B7, B8, B9, B11) were used for scATAC-seq. The snATAC-seq library construction is done for these 8 hippocampal samples using Chromium Single Cell ATAC Library & Gel Bead Kit v2 chemistry.

### Single-cell ATAC-seq library processing and alignment

snATAC-seq libraries were generated and processed using the Cell Ranger ATAC pipeline (v2.1.0, 10x Genomics). Raw reads from 8 hippocampal samples were aligned to the mm10 reference genome (refdata-cellranger-arc-mm10-2020-A-2.0.0). Peak-barcode matrices and fragment files were generated using cellranger-atac count and used for downstream analysis.

### Unified-peak set workflow

Downstream analysis was performed in R using the Signac (v1.13.0) and Seurat (v5.1.0) packages. Individual peak files were merged into a unified peak set using reduce() from the GenomicRanges package, excluding peaks <20 bp or >10,000 bp. Cells with fewer than 500 fragments in peaks were filtered out. Chromatin assays were constructed for each sample using FeatureMatrix().

Each Seurat object was annotated using EnsDb.Mmusculus.v79, and quality control (QC) metrics were computed: transcription start site (TSS) enrichment, nucleosome signal, percent reads in peaks, and blacklist ratio (based on blacklist_mm10). Cells were filtered using quantile-based thresholds, excluding outliers (below 2nd or above the 98th percentile for key metrics) to remove low-quality cells and potential multiplets.

Filtered objects were merged, then the merged object was processed with normalization with RunTFIDF(), dimensionality reduction via latent semantic indexing (RunSVD()), batch correction with harmony (RunHarmony() using LSI dims 2–30), and UMAP embedding. Clusters were identified with FindNeighbors() and FindClusters() with resolution 0.8 and algorithm 1. Gene activity scores were calculated via GeneActivity() and log-normalized with NormalizeData() (scale factor = median of nCount_Activity). A preprocessed single-nucleus RNA-seq dataset (Singh et al., 2023) served as a reference for cell type annotation. Unmatching samples (A3, A5, B10, B12) were excluded. Anchors were computed using canonical correlation analysis (CCA) to predict cells in scATAC-seq dataset.

### Sample-specific peak set workflow

In addition to the unified peak set approach, we conducted a complementary analysis using sample-specific peak sets generated by Cell Ranger ATAC. For each sample, peak-barcode matrices and fragment files were independently loaded, and chromatin assays were constructed using a minimum cell threshold of min.cells =10. Seurat objects were generated separately for each sample and subsequently merged for integrated downstream analysis. Gene annotations were added using the EnsDb.Mmusculus.v79 database. QC metrics were calculated for each cell, including TSS enrichment, nucleosome signal, percent reads in peaks, and the blacklist ratio. To ensure high-quality samples, cells were filtered using the following strict criteria: nCount_peaks between 3,000 and 30,000, pct_reads_in_peaks >15%, blacklist_ratio < 0.05, nucleosome_signal <4, and TSS.enrichment >3.

Dimensionality reduction and clustering were performed: normalization with RunTFIDF(), feature selection via FindTopFeatures(min.cutoff =’q0’), latent semantic indexing (LSI) using RunSVD() on dimensions 2 through 8, and UMAP embedding via RunUMAP(). Clusters were identified using FindNeighbors() and FindClusters() with algorithm 1. Gene activity scores were computed using GeneActivity() and log-normalized with NormalizeData(). Cell type annotations previously inferred from the unified peak set analysis were transferred to the sample-specific peak set dataset by matching shared cell barcodes. Cells without annotation were excluded from further analysis.

For analysis related to the Pyramidal cluster, pyramidal neurons were subset from the unified peak set dataset and subjected to reclustering to refine subtype resolution. To facilitate annotation, these cells were integrated with a previously annotated single-nucleus RNA-seq reference dataset (Singh et al., 2023) using CCA, with 30 dimensions and 10 nearest neighbors employed to identify cross-modality anchors. Uncharacterized or ambiguous Seurat clusters were either removed or reannotated based on their transcriptomic profiles and proximity to well-defined clusters in the reference dataset. The finalized annotations were then transferred to the corresponding pyramidal neuron populations within the sample-specific peak set dataset by matching shared cell barcodes.

### Differential Accessibility (DA) Analysis

All rhEPO- and placebo-derived cells were combined within each cluster (seurat cluster, higher-level cell-type annotation, or pyramidal subclusters). We then executed FindMarkers (test.use = “LR”, min.pct = 0.01, latent.vars = “nCount_peaks”) in a one-versus-rest fashion to identify peaks that are intrinsically enriched or depleted in a given cluster relative to all other clusters. We used 0.01 as min.pct threshold to capture accessibility shifts in smaller subfractions of a population. Peaks passing FDR < 0.01 were considered cell-type DA peaks. Furthermore, to pinpoint treatment-dependent changes, we stratified cells by condition inside each cluster and ran FindMarkers() with min.pct = 0.01 in the LR framework, and depth correction was applied. Peaks with FDR < 0.01 were classified as condition DA peaks. In differences between defined clusters (cell-type DA peaks) or treatment-dependent changes (condition DA peaks), summits (peak mid-points) were annotated via TxDb.Mmusculus.UCSC.mm10.knownGene to the closest gene’s promoter, binned by genomic context (promoter, exon, intron, intergenic) and intersected with RepeatMasker (mm10) to identify and quantify overlap with TEs. Low-complexity, satellite, RNA, rRNA, tRNA, scRNA, snRNA, RC, srpRNA, and ambiguous repeats (entries flagged with “?” such as “LINE?”) were excluded from all TE analyses unless explicitly stated.

### Cluster-resolved motif enrichment

Chromatin dynamics were analyzed in two complementary tiers: chromatin accessibility differences between defined clusters and treatment-dependent changes. Vertebrate position-frequency matrices from the JASPAR-2020 CORE collection were added for motif analysis. For every Seurat cluster, higher-level cell-type annotation, or pyramidal subclusters, identified DA peaks using FindMarkers were filtered as min.pct threshold is 0.10, only positive, and FDR < 0.05. Clusters contributing ≥ 10 significant peaks were subjected to motif enrichment with FindMotifs(). Likewise, within the same clustering framework, cells were split by condition, and DA peaks filtered with min.pct threshold is 0.10, only positive, and FDR < 0.05 parameters were used for motif analysis. For any cluster yielding ≥ 10 rhEPO-responsive peaks, FindMotifs() was run to identify transcription-factor motifs enriched in rhEPO-gained chromatin.

### Transcription-factor ChIP-seq integration

Genome-wide binding profiles for 13 neurodevelopmental transcription factors (ASCL1, EGR1, FOS, FOXG1, FOXO3, JUN, NEUROD1, NEUROD2, NEUROG2, PAX6, SOX2, TBR1) were downloaded from GEO, (Table S17). Peak files were lifted over to the mm10 assembly where necessary (UCSC liftOver) and collapsed into non-redundant consensus regions with bedtools merge function. To ask whether individual TFs preferentially occupy repeat-derived regulatory DNA, we quantified intersections between each consensus ChIP peak set and RepeatMasker annotations using regioneR; 1000 permutations provided empirical P values and observed/expected overlap ratios. TE-associated peaks were assigned to the nearest transcription-start site with ChIPseeker (TxDb.Mmusculus.UCSC.mm10.knownGene) and subjected to GO-biological-process enrichment with clusterProfiler::enrichGO (BH-adjusted q ≤ 0.01). Next, TF peaks were overlapped with the 11 broad cell-class DA peak lists and the refined pyramidal-lineage DA peaks identified in our snATAC-seq analysis.

### Data availability

All raw FASTQ files, fragment files, count matrices, and intermediate Seurat objects underlying the present study have been deposited in the GEO GSE and zenodo (available upon a reasonable request).

### Code availability

Code for data processing, quality control, clustering, differential-accessibility testing, motif analysis, and TF-ChIP overlap is available on the following GitHub link: https://github.com/umutcakir/ATAC_EPO_vs_Placebo.

## Acknowledgements

This work has been supported by the European Research Council (ERC) Advanced Grant to HE under the European Union’s Horizon Europe research and innovation programme (acronym *BREPOCI*; grant agreement No 101054369), as well as by the Max Planck Society, the Max Planck Förderstiftung, the Deutsche Forschungsgemeinschaft (DFG, German Research Foundation) via DFG-Center for Nanoscale Microscopy & Molecular Physiology of the Brain (CNMPB). Research in the labs of HE and KAN is funded by SFB TRR 274/2 – 408885537 project C01 (to H.E. and K.A.N). UC received support from the IMPRS-Genome Science PhD program. Research in the labs of DG, RK and KAN is supported by the Adelson Medical Research Foundation.

## Author Contributions

Concept, design, supervision: MS, HE

Funding acquisition: HE, RK, DG, KAN

Drafting manuscript and display items: UC, MS, MF, UJB, HE

Data acquisition/generation: MS, RK, DG, KAN

Data analyses & interpretation: UC, RK, MS, HE

**All authors read and approved the final version of the manuscript.**

## EXTENDED DATA

**Extended Data Fig. S1.**
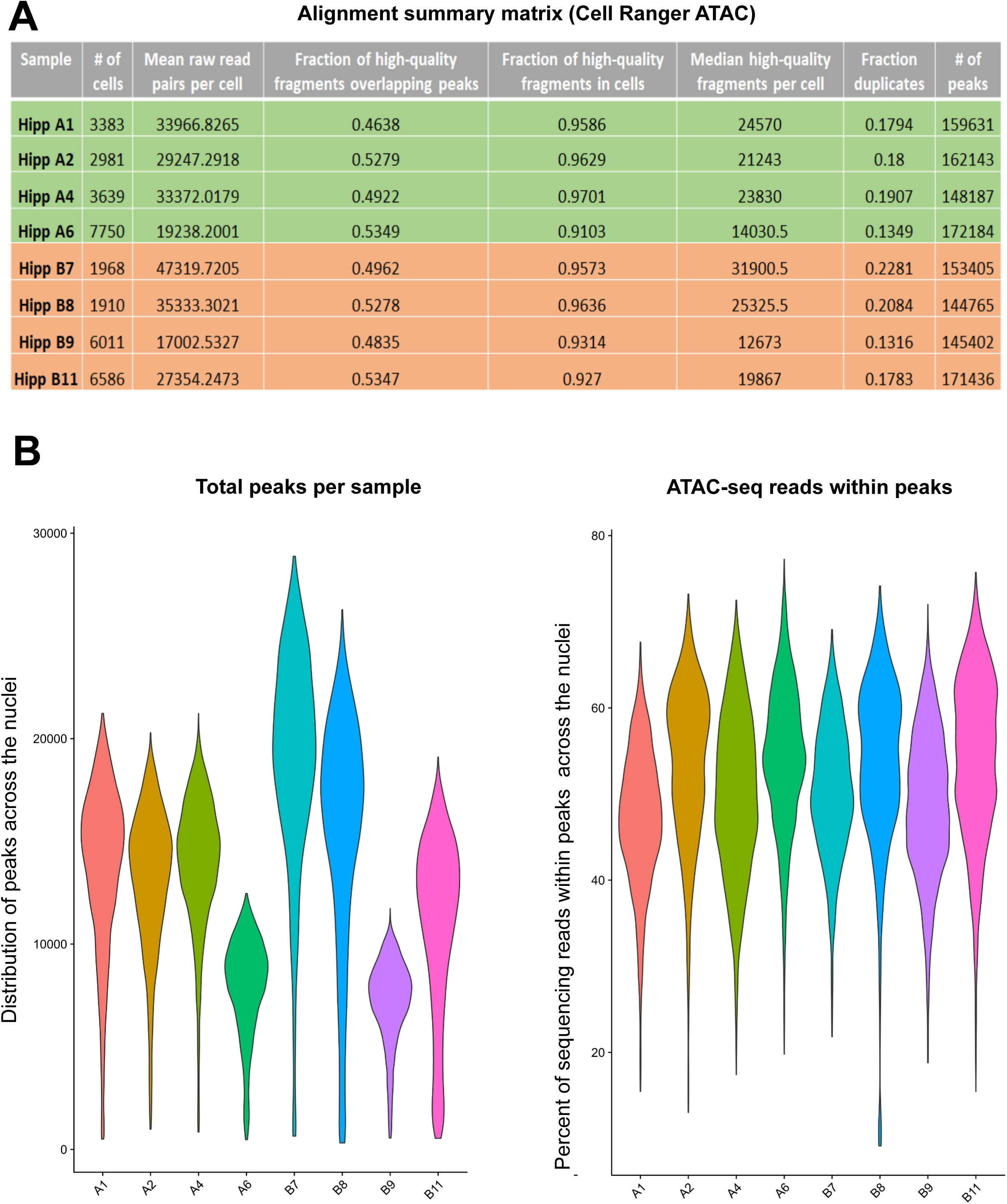
Quality control metrics derived from Cell Ranger ATAC output. **(A)** Summary of snATAC-seq quality metrics obtained from Cell Ranger ATAC output. For each sample, the table shows the number of high-quality nuclei passing filtering, mean raw read pairs per nucleus, fraction of high-quality fragments overlapping called peaks, fraction of high-quality fragments confidently assigned to nuclei, median high-quality fragments per nucleus, duplicate fragment fraction, and total number of peaks detected. **(B)** Violin plots showing (left) the distribution of total peaks detected per nucleus and (right) the percentage of sequencing reads falling within peaks across nuclei for each sample.

**Extended Data Fig. S2.**
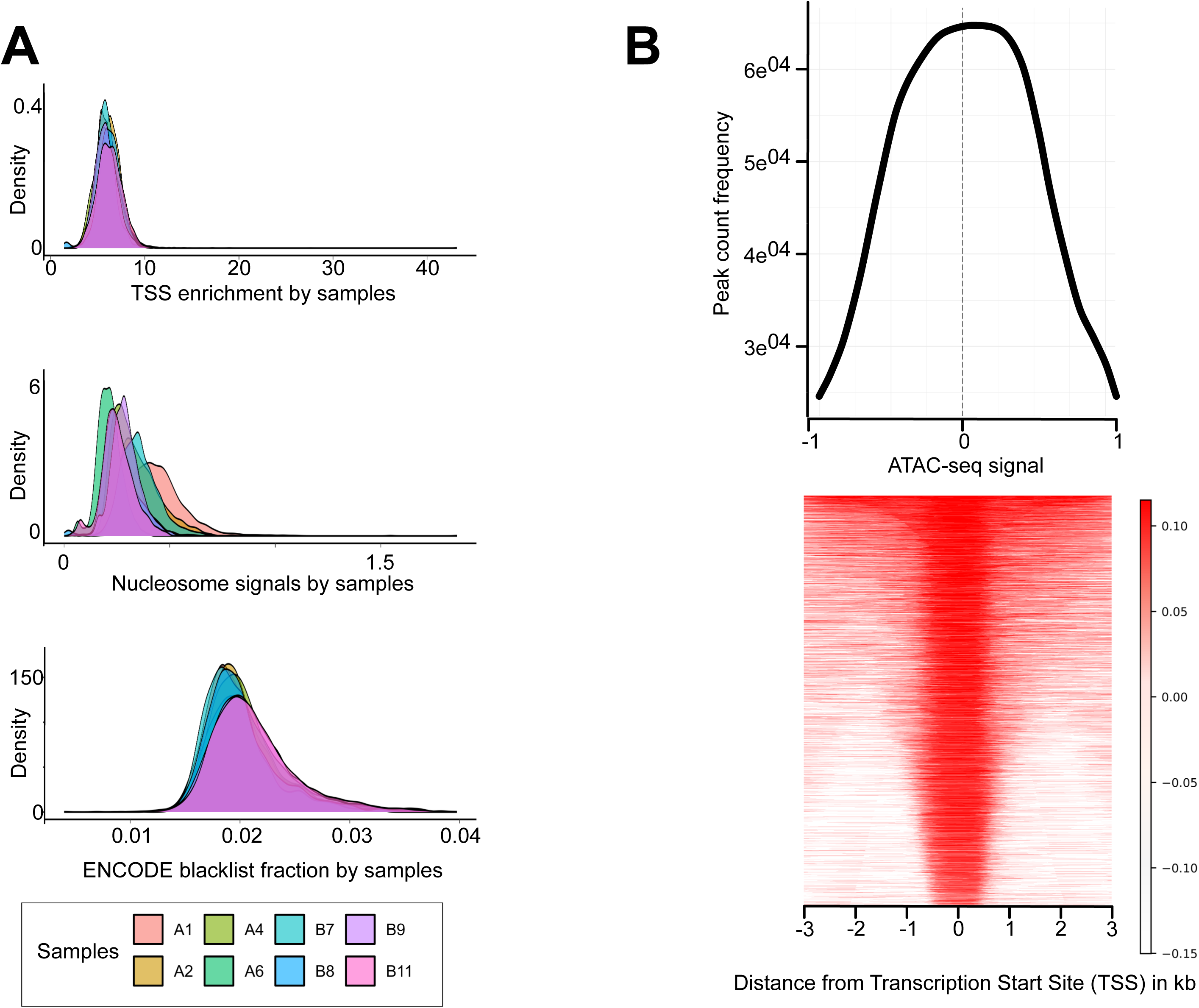
Additional quality control profiles for snATAC-seq libraries. **(A)** Density distributions of three core snATAC-seq quality metrics across samples: TSS enrichment scores, nucleosome signal estimates, and ENCODE blacklist fractions. TSS enrichment reflects the characteristic accumulation of fragments at transcription start sites, while nucleosome signal is the ratio of mononucleosome fragments to nucleosome-free fragments per cell. ENCODE blacklist fractions indicate the proportion of fragments mapping to problematic genomic regions. **(B)** Aggregate ATAC-seq signal profile centered on transcription start sites (top), showing the expected sharp increase in accessibility at TSSs. The heatmap (bottom) displays ATAC-seq fragment density within ±3 kb of TSSs for all peaks, illustrating strong signal localization around promoter regions and consistent accessibility patterns across the dataset.

**Extended Data Fig. S3.**
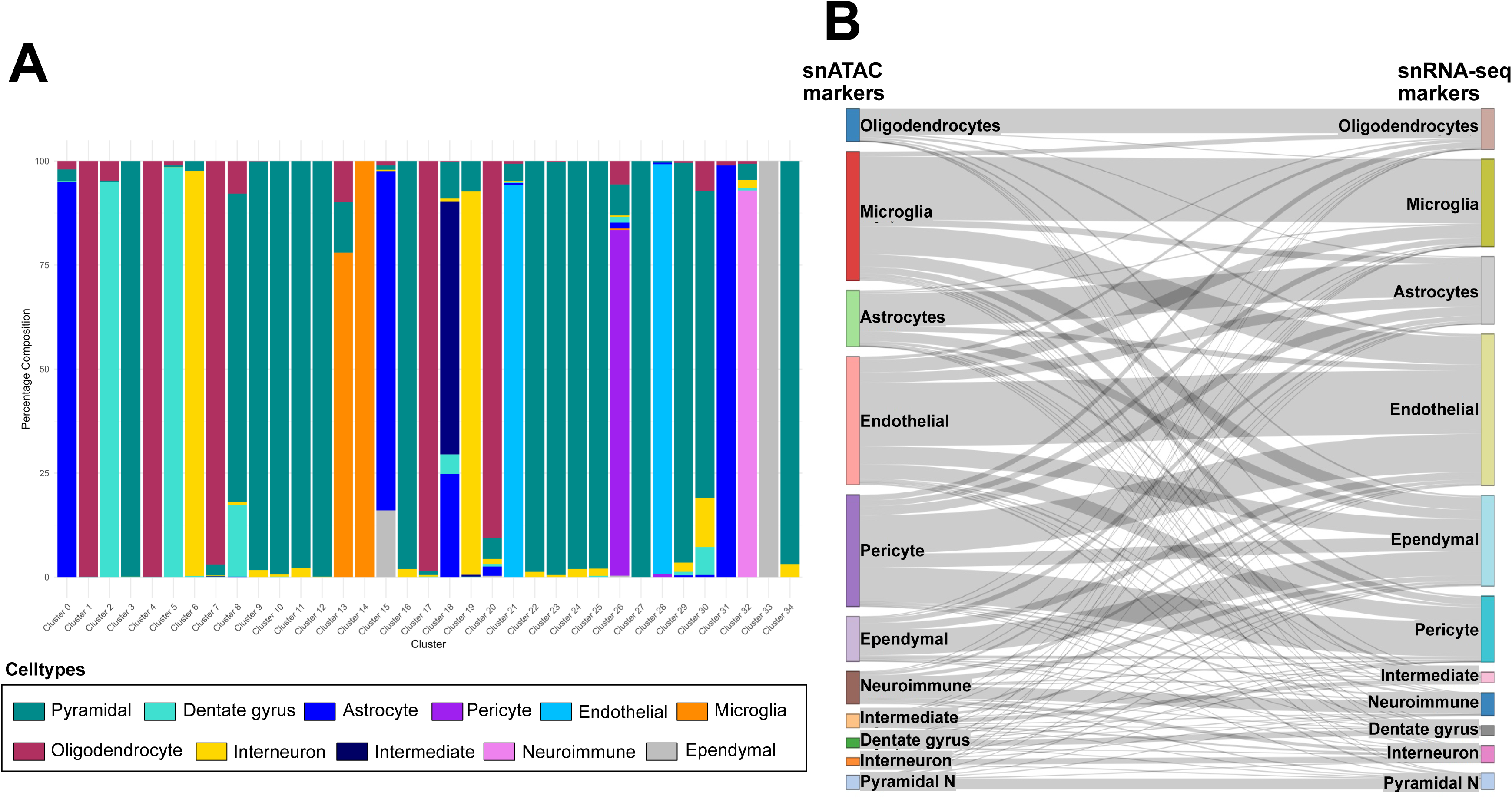
Cross-modality cell-type annotation and correspondence between snATAC-seq and snRNA-seq datasets. **(A)** Stacked bar plot showing the percentage composition of annotated cell types within each Seurat snATAC-seq cluster. Each bar represents one chromatin-accessibility-defined Seurat cluster, and colors correspond to major hippocampal lineages: pyramidal neurons, dentate gyrus neurons, astrocytes, oligodendrocytes, interneurons, microglia, endothelial cells, pericytes, intermediate cells, neuroimmune cells, and ependymal cells. The distribution illustrates the dominant cell identity assigned to each Seurat snATAC-seq cluster. **(B)** Sankey diagram comparing cell-type annotations derived from snATAC-seq markers (left) with those obtained from snRNA-seq markers (right). Flow widths represent the degree of correspondence between modalities, with thicker connections indicating stronger agreement between ATAC-based and RNA-based cell-type assignments.

**Extended Data Fig. S4.**
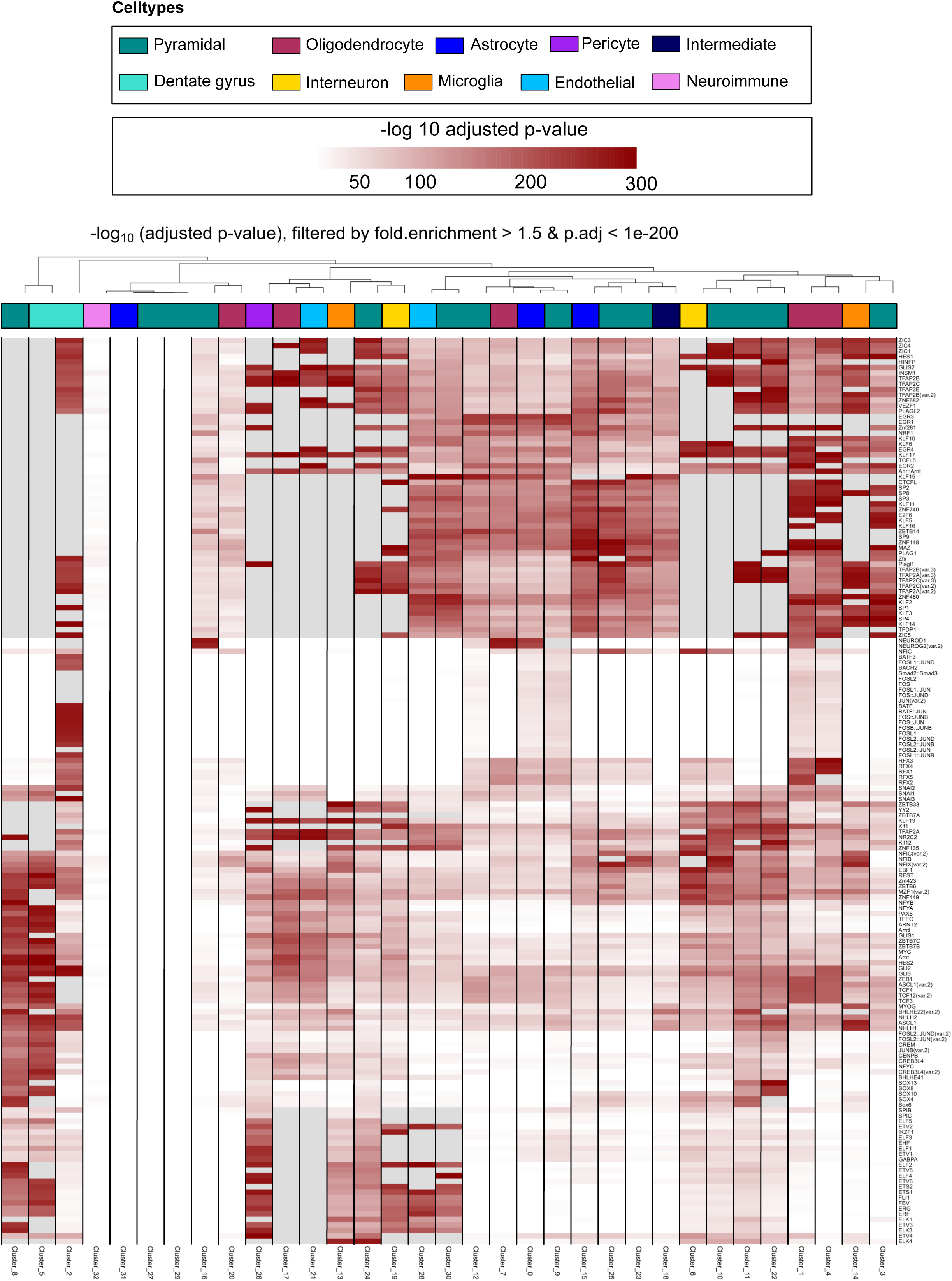
Transcription factor motif enrichment across Seurat snATAC-seq clusters. Heatmap showing enriched transcription factor motifs across Seurat-defined snATAC-seq clusters. Motifs were filtered for fold enrichment > 1.5 and adjusted p-value < 1×10⁻²_J_J. Each column represents an individual Seurat cluster and is annotated by its corresponding cell-type identity (top color bar). Each row shows a transcription factor motif, with color intensity indicating the −log₁₀ adjusted p-value of enrichment. The pattern of enrichment reveals distinct regulatory signatures across clusters, with related cell types sharing similar motif profiles and exhibiting coordinated accessibility at lineage-specific regulatory elements.

**Extended Data Fig. S5.**
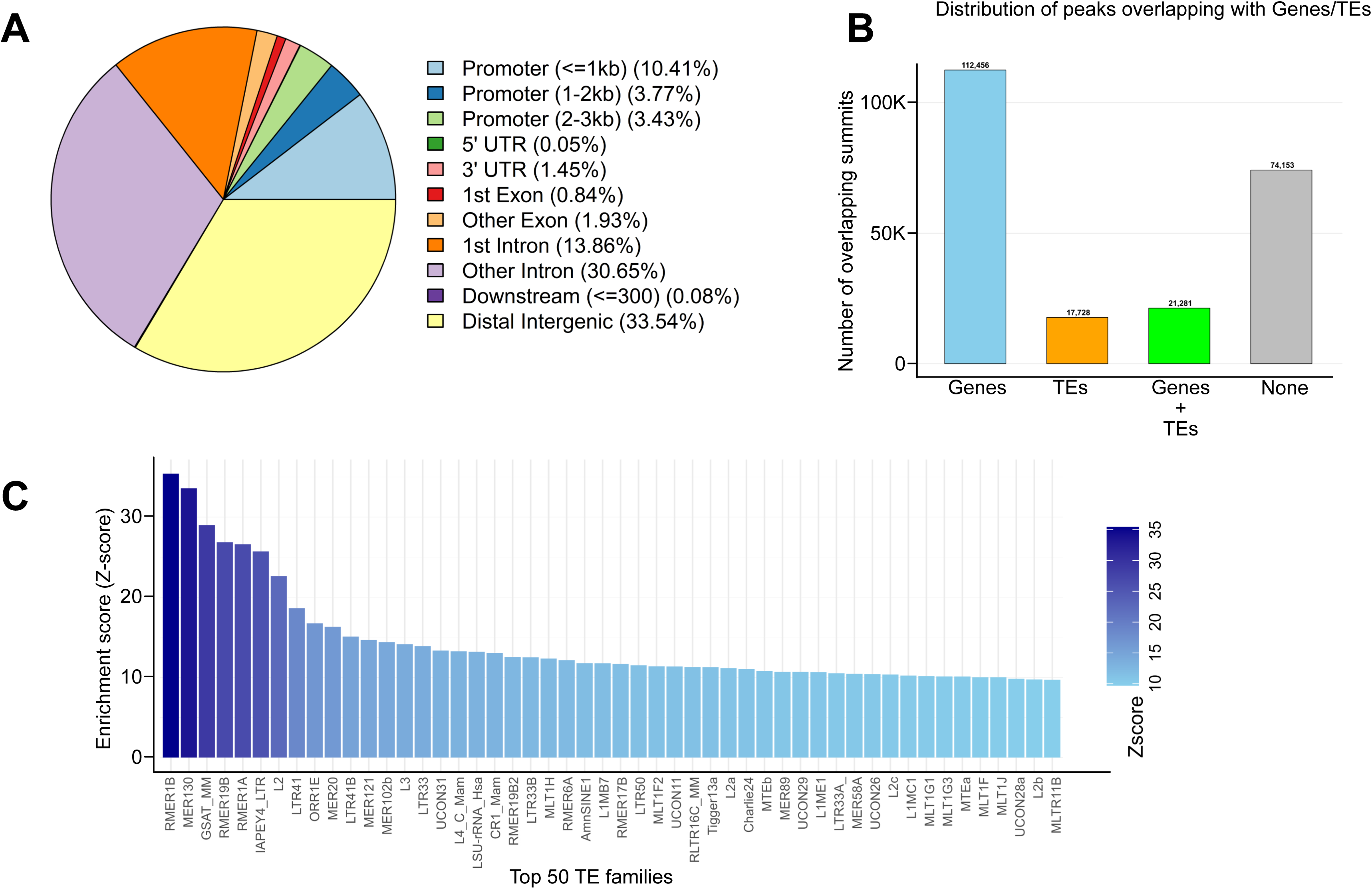
Genomic distribution of snATAC-seq peaks and enrichment of transposable element families. **(A)** Pie chart summarizing the genomic annotation of all snATAC-seq peak summits, showing the proportion of peaks located in promoter regions (stratified by distance to the TSS), untranslated regions, exons, introns, downstream regions, and distal intergenic regions. Distal intergenic and intronic regions constitute the largest fractions of accessible chromatin. **(B)** Bar plot showing the number of peak summits overlapping annotated genes, annotated transposable elements (TEs), both genes and TEs, or neither genomic feature. A substantial subset of peaks overlaps TEs, either exclusively or in combination with gene bodies. **(C)** Z-scores for the enrichment of the top 50 TE families overlapping snATAC-seq peak summits. TE families are ranked by their enrichment, revealing pronounced overrepresentation of several LTR, SINE, and LINE subfamilies within accessible chromatin regions.

**Extended Data Fig. S6.**
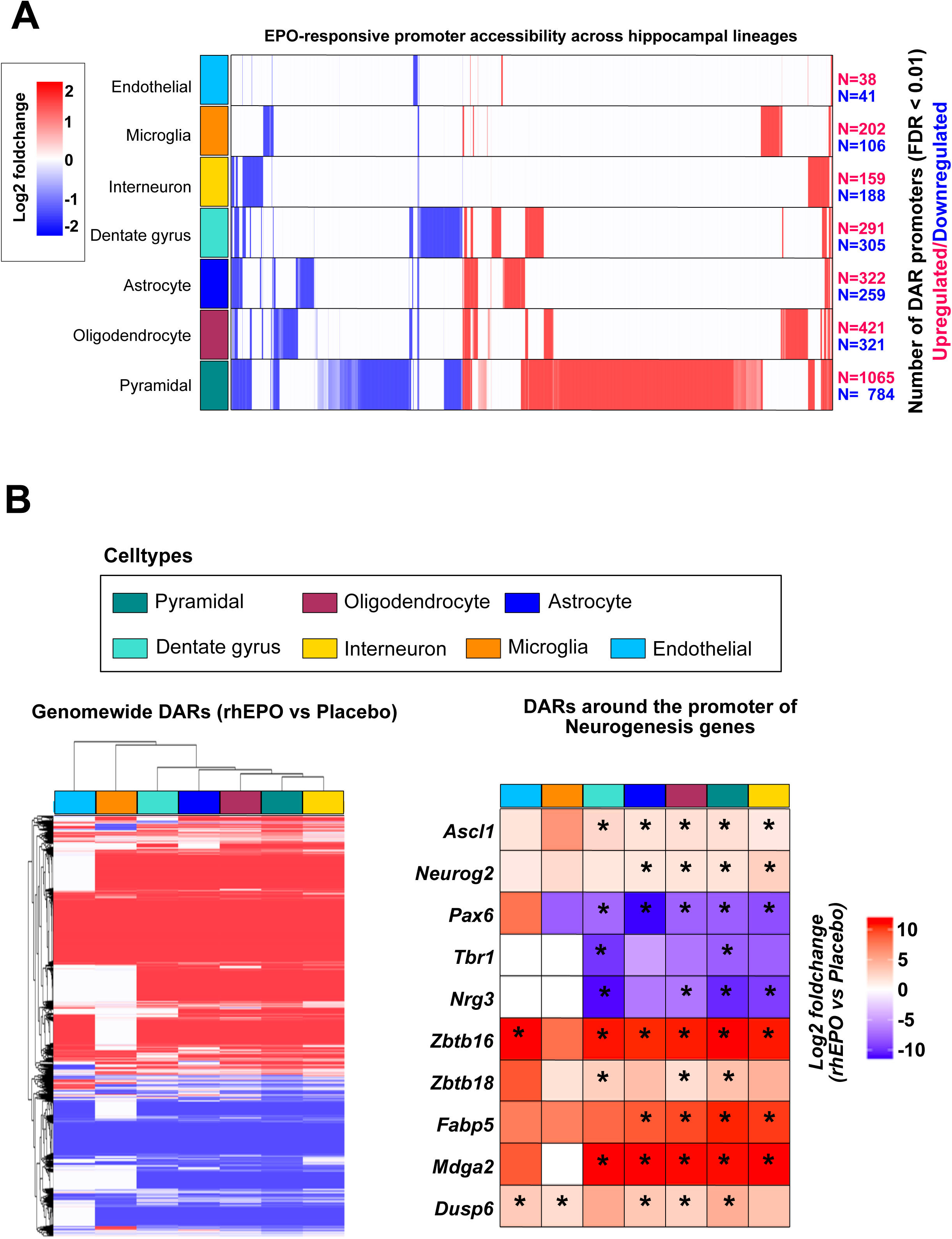
EPO-responsive promoter accessibility across hippocampal lineages. **(A)** Heatmap showing log_2_ fold-change in promoter accessibility (FDR < 0.01) between rhEPO-and placebo-treated samples across major hippocampal cell types. Each row corresponds to a lineage and each column to a promoter-associated differentially accessible region (DAR). Red indicates increased accessibility under rhEPO, and blue indicates decreased accessibility. The total number of upregulated and downregulated promoter DARs for each lineage is shown on the right. Pyramidal neurons exhibit the largest number of EPO-responsive promoters. **(B)** Differential accessibility heatmaps summarizing genome-wide DARs (left) and promoter-specific DARs for selected neurogenesis-related genes (right) across hippocampal cell types. The genome-wide heatmap depicts hierarchical clustering of all DARs between rhEPO and placebo, with colors indicating log_2_ fold-changes. The promoter-focused heatmap highlights accessibility changes at key neurogenic regulators; these genes are Ascl1, Neurog2, Pax6, Tbr1, Nrg3, Zbtb16, Zbtb18, Fabp5, Mdga2, and Dusp6. Asterisks denote significant change upon the rhEPO-treatment (FDR < 0.01).

**Extended Data Fig. S7.**
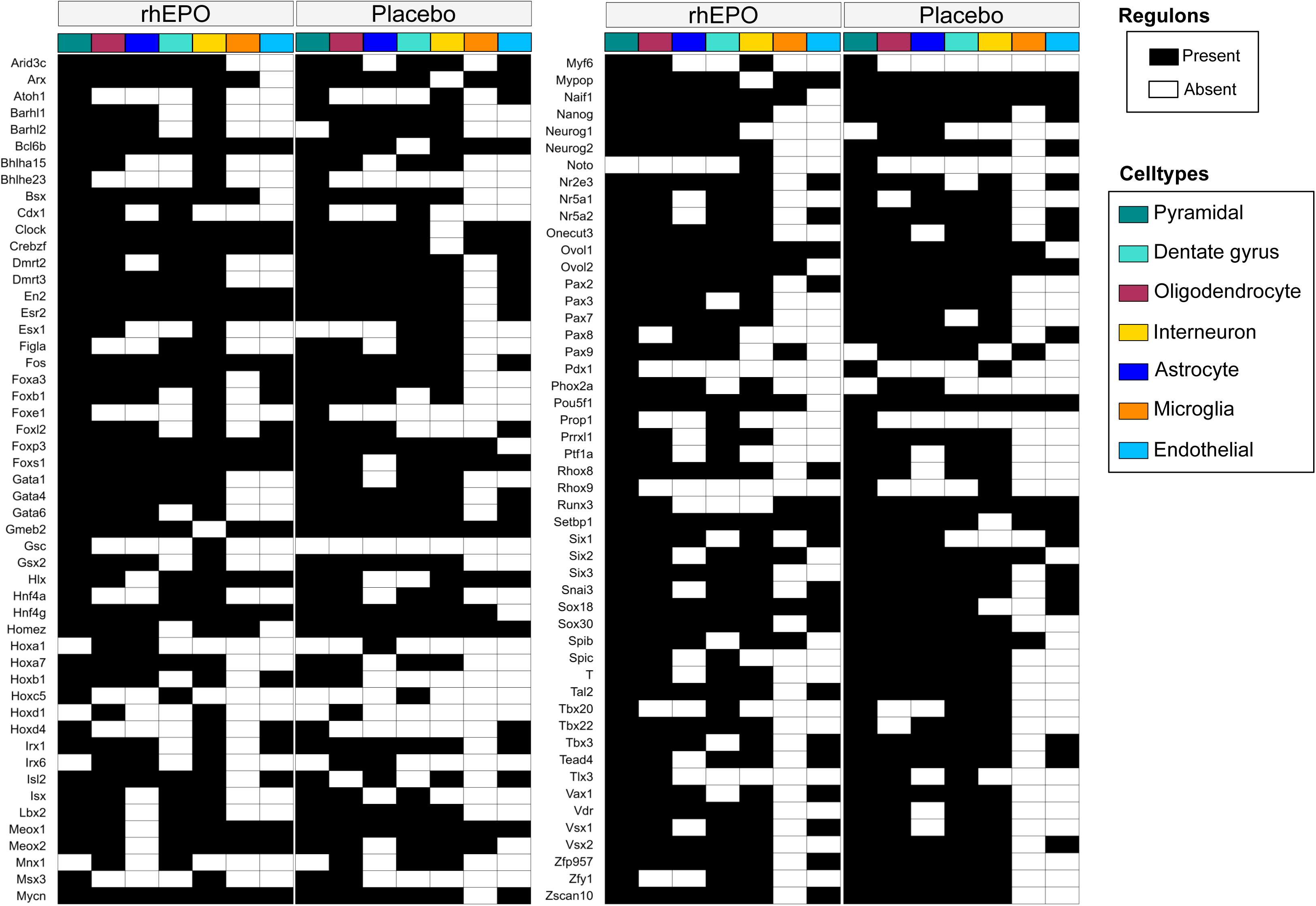
Regulon activity across hippocampal lineages under rhEPO and placebo conditions. Binary heatmap showing the presence or absence of Pando-inferred regulons across major hippocampal cell types for rhEPO-treated and placebo-treated conditions. Each row represents a regulon, and each column corresponds to a cell type, annotated with color-coded lineage identities. Black filled boxes indicate regulons that are active (present) in a given lineage, while white filled boxes indicate the absence of regulon activity. The comparison highlights regulons that are active or absent under rhEPO or placebo treatment and reveals lineage-specific differences in regulatory program activation across the hippocampus.

**Extended Data Fig. S8.**
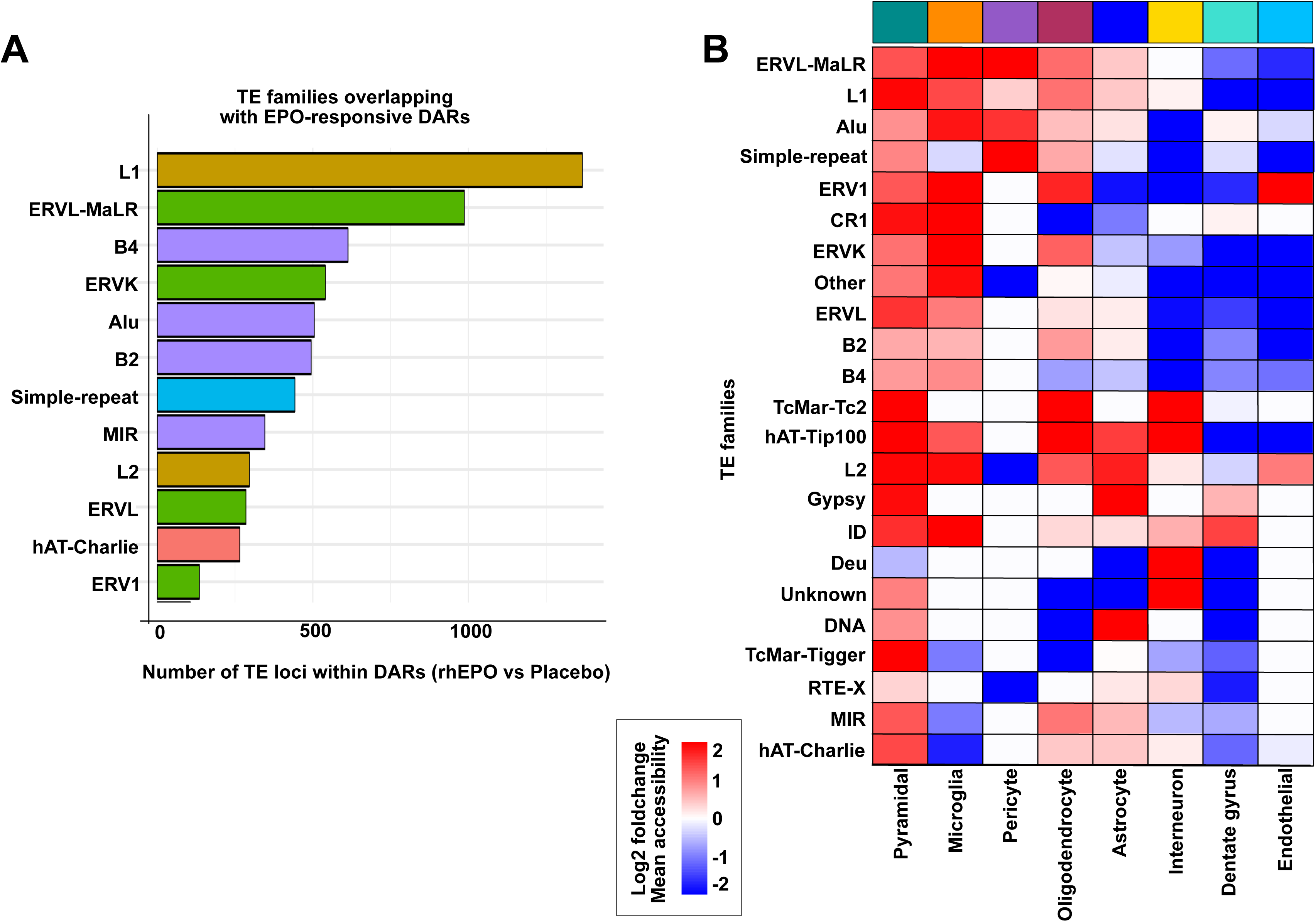
Transposable element families enriched within EPO-responsive chromatin regions. **(A)** Bar plot showing the number of transposable element (TE) loci overlapping differentially accessible regions (DARs) identified between rhEPO- and placebo-treated samples. TE families are ranked by the number of overlapping loci, revealing strong contributions from L1, ERVL-MaLR, B4, ERVK, Alu, B2, and other repeat classes to EPO-responsive changes in chromatin accessibility. **(B)** Heatmap displaying the log_2_ fold-change in accessibility for TE-associated DARs across major hippocampal cell types. Rows represent TE families, and columns represent cell lineages. Color intensity reflects mean chromatin accessibility differences (rhEPO vs. placebo), with red indicating increased accessibility and blue indicating decreased accessibility. The heatmap highlights lineage-specific patterns of TE-derived accessibility remodeling in response to rhEPO treatment.

**Extended Data Fig. S9.**
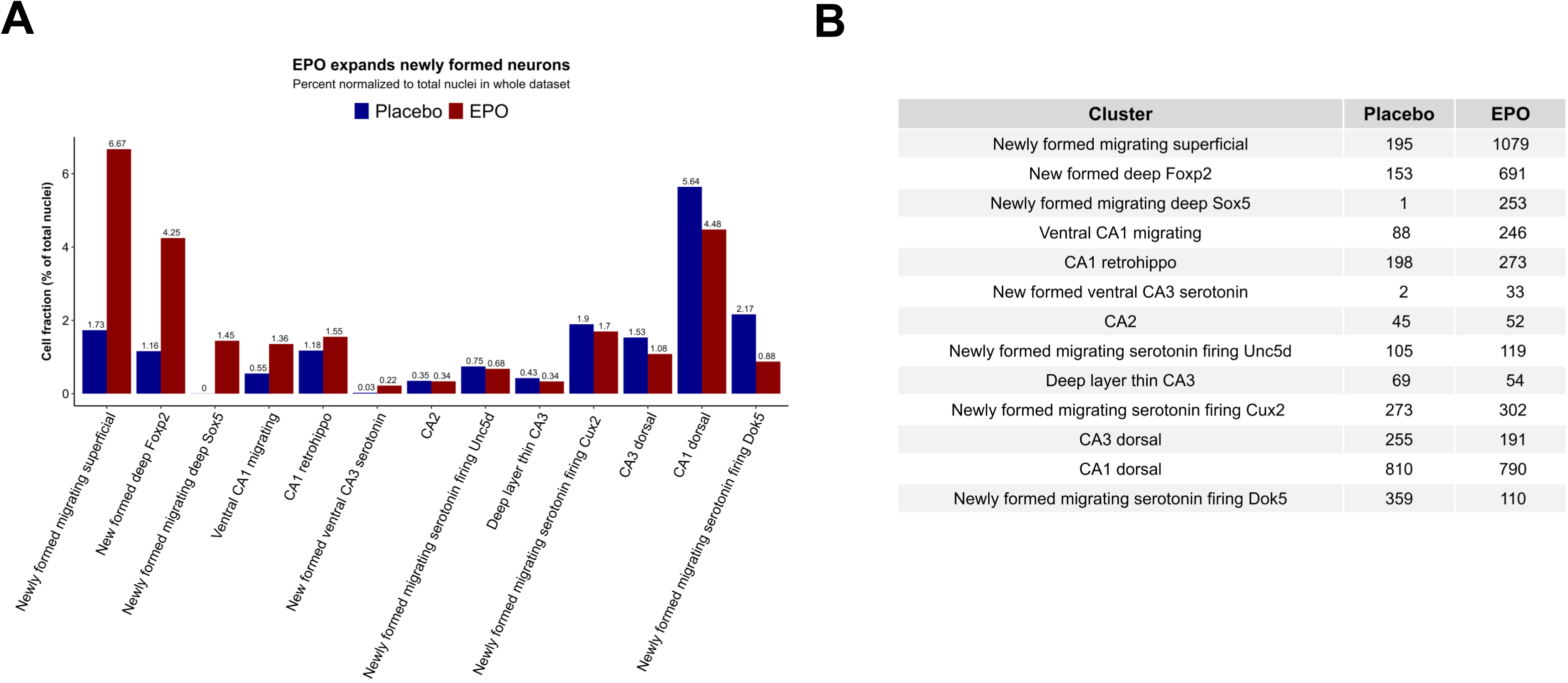
Expansion of newly formed pyramidal neuron populations under rhEPO treatment. **(A)** Bar plot showing the relative fraction of newly formed pyramidal neuron subtypes in placebo- and rhEPO-treated hippocampi, normalized to the total number of nuclei. Across multiple early-stage neuronal populations, including superficial migrating, deep-layer Foxp2⁺ and migrating deep Sox5⁺ neurons, ventral CA1 migrating, CA1 retrohippocampal, and ventral CA3 serotonin neuron subtypes, rhEPO treatment leads to a substantial increase in cell abundance compared with placebo conditions. **(B)** Table summarizing the absolute number of nuclei assigned to each newly formed pyramidal neuron cluster under placebo and rhEPO conditions. The numerical increases further illustrate the strong expansion of newly formed pyramidal lineages following rhEPO treatment.

**Extended Data Fig. S10.**
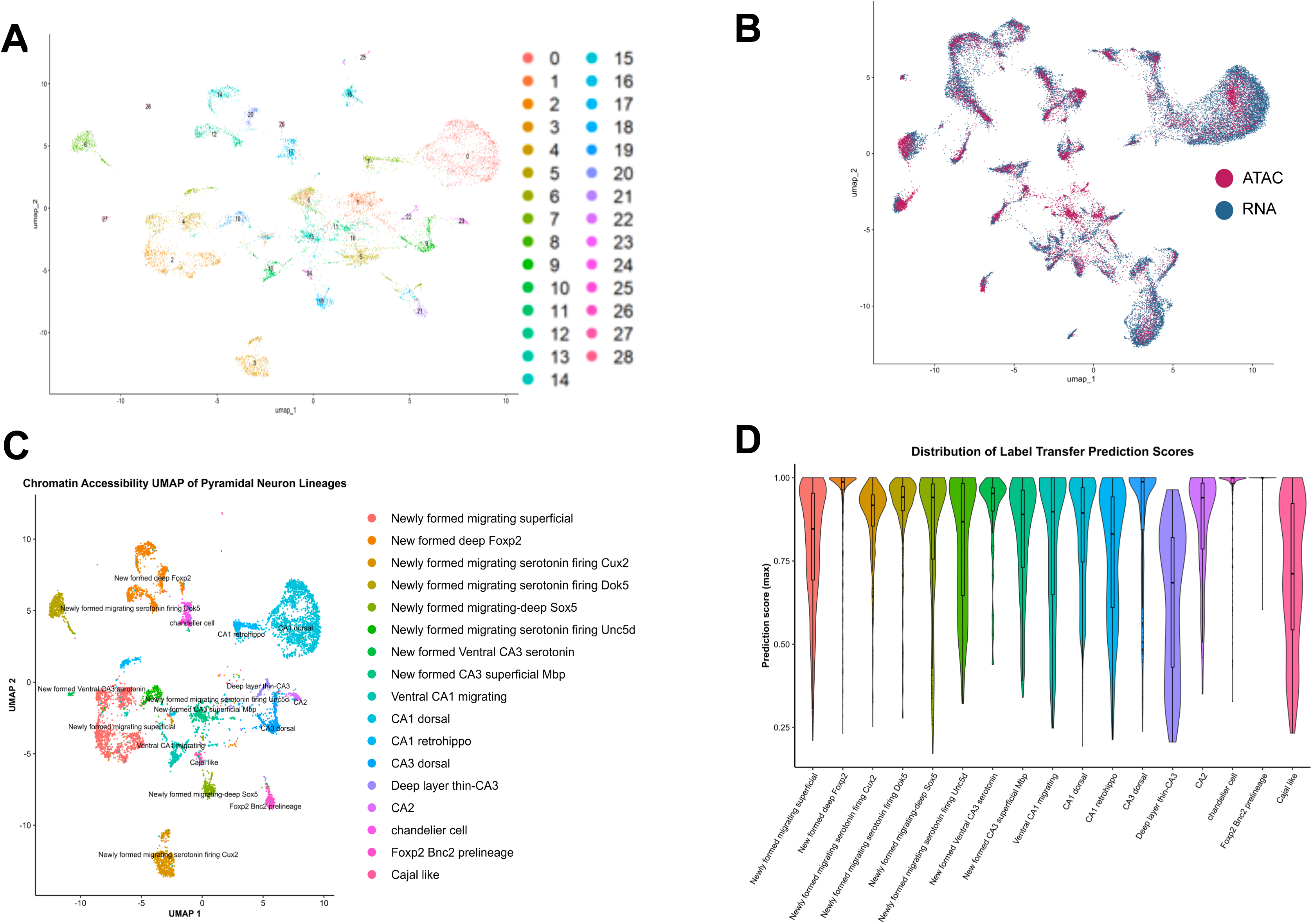
Integration of snATAC-seq and snRNA-seq datasets and annotation of pyramidal neuron lineages. **(A)** UMAP representation of Seurat-defined chromatin accessibility clusters from snATAC-seq, with each color indicating a distinct cluster ID. **(B)** Joint UMAP embedding of snATAC-seq (red) and snRNA-seq (blue) datasets, showing strong overlap between modalities following integration and confirming successful alignment of chromatin accessibility and transcriptomic profiles. **(C)** UMAP visualization of chromatin accessibility profiles for annotated pyramidal neuron sublineages. **(D)** Violin plots of label transfer prediction scores for pyramidal neuron subtypes, reflecting the confidence of cross-modality cell identity assignments.

**Extended Data Fig. S11.**
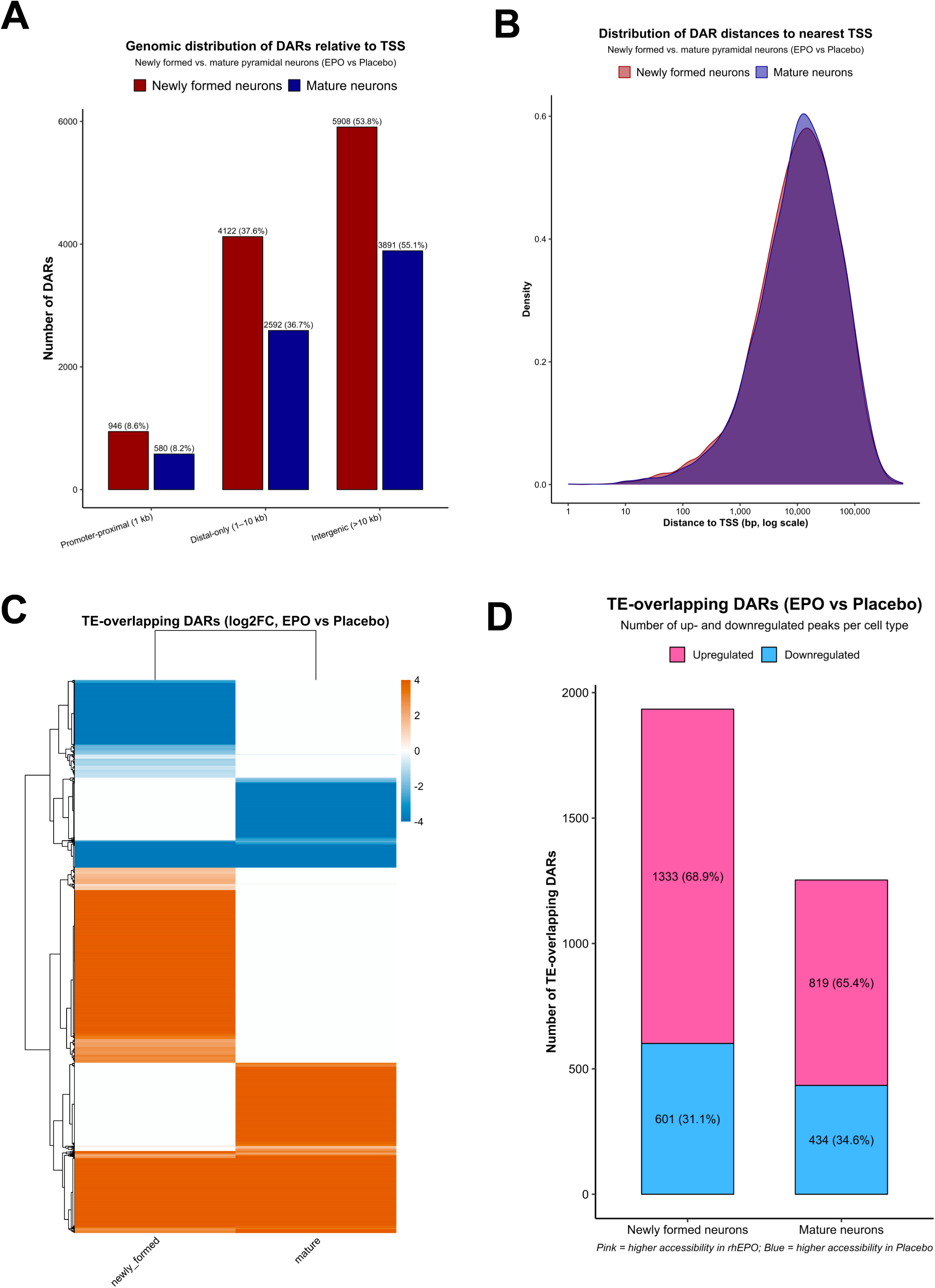
Genomic distribution and Transposable element (TE) association of differentially accessible regions (DARs) in newly formed and mature pyramidal neurons. **(A)** Bar plot showing the number of DARs located in promoter-proximal regions (≤1 kb), distal regions (1–10 kb), and intergenic regions (>10 kb) for newly formed and mature pyramidal neurons (rhEPO vs. placebo). Newly formed neurons exhibit a larger number and higher proportion of distal and intergenic DARs. **(B)** Density plot of distances from DAR summits to the nearest transcription start site (TSS) on a log-scaled x-axis. Newly formed and mature neurons show broadly similar distance profiles, with both distributions peaking around 10 kilobases from the nearest TSS **(C)** Heatmap of log_2_ fold-change values for TE-overlapping DARs in newly formed and mature neurons (rhEPO vs. placebo). Rows represent TE-associated peaks, illustrating distinct groups of TE-derived regions that become more or less accessible under EPO treatment depending on neuronal maturation state. **(D)** Bar plot summarizing the number of TE-overlapping DARs that increase (pink) or decrease (blue) in accessibility in response to rhEPO within newly formed and mature neurons. Newly formed neurons exhibit a greater number of rhEPO-induced accessibility gains or losses compared to mature neurons.

**Extended Data Fig. S12.**
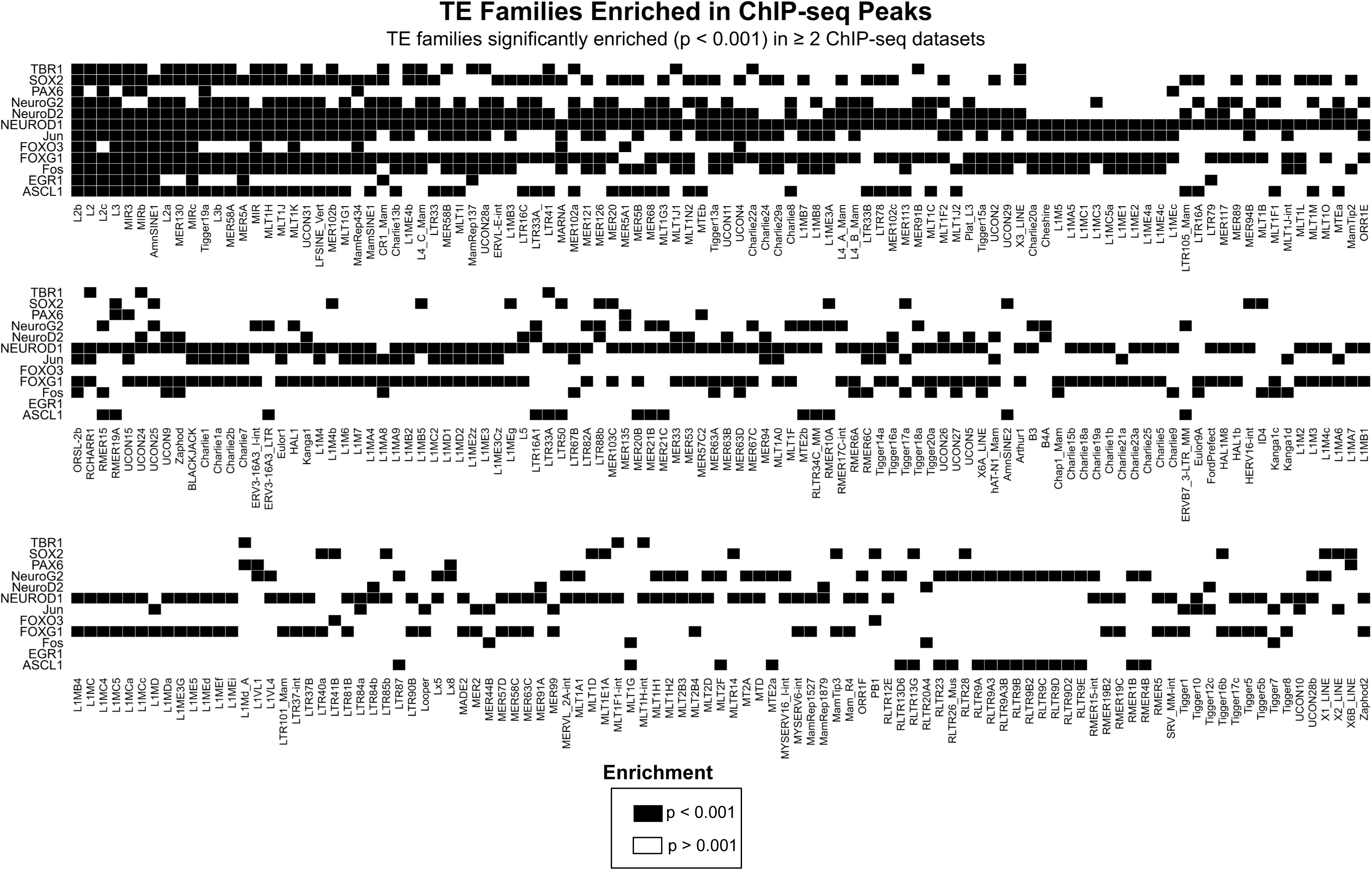
Transposable element families enriched within neurogenic transcription factor ChIP-seq peaks. Matrix showing TE families significantly enriched (p < 0.001) within ChIP-seq binding sites of neurogenic transcription factors. Rows correspond to transcription factors (TBR1, SOX2, PAX6, NEUROG2, NEUROD2, NEUROD1, JUN, FOS, FOXO3, FOXG1, EGR1, ASCL1), and columns represent individual TE families. Black squares indicate significant enrichment (p < 0.001), while white squares indicate non-significant enrichment (p > 0.001). Only TE families enriched in ≥2 independent ChIP-seq datasets are shown. The matrix highlights recurrent engagement of diverse TE families by key neurogenic TFs, supporting widespread co-option of TE-derived sequences in neuronal regulatory networks.

**Extended Data Fig. S13.**
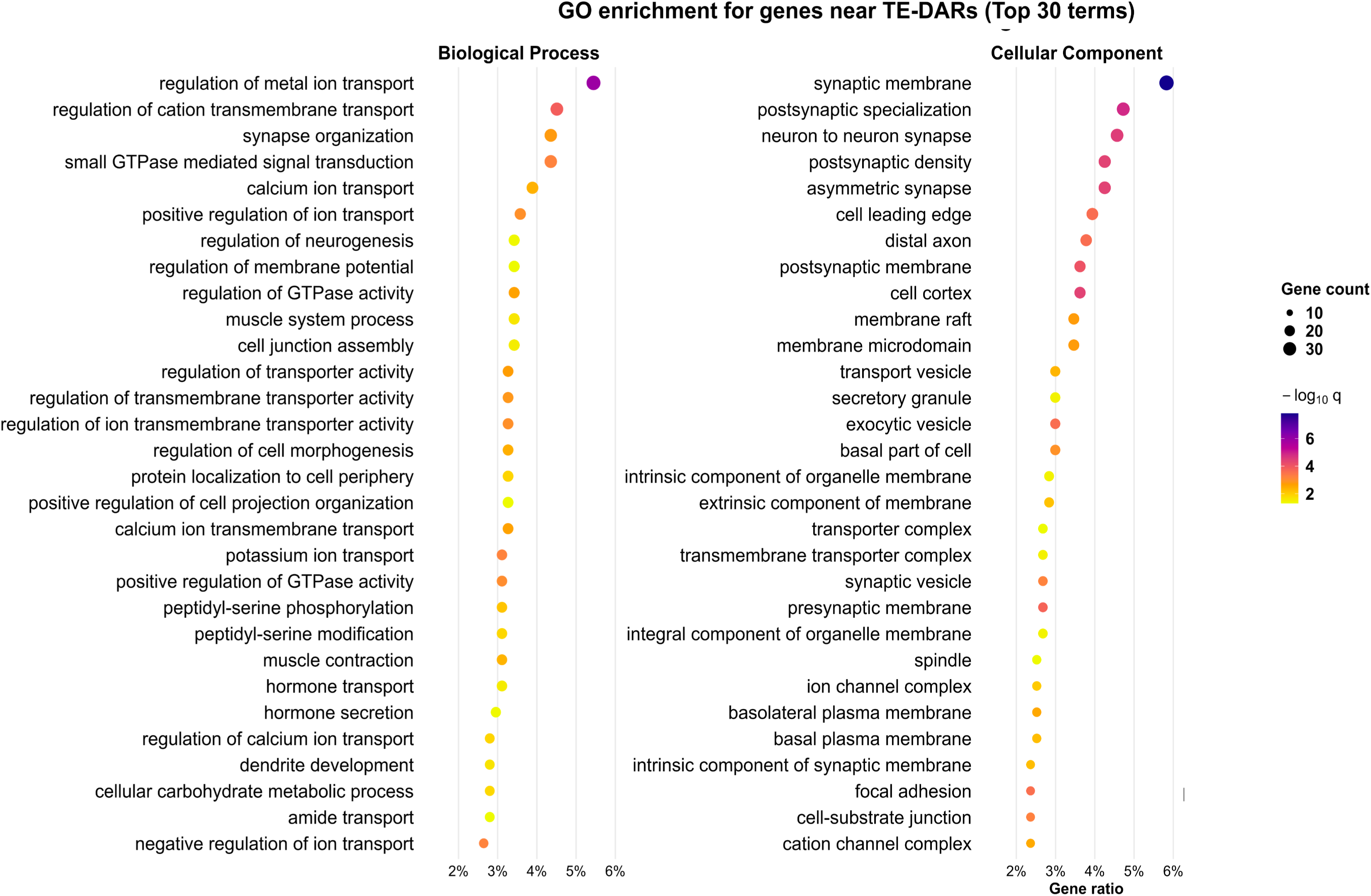
Gene ontology enrichment for genes located within ±10 kb of TE-overlapping DARs. Dot plots showing the top 30 enriched Gene Ontology (GO) terms for genes whose transcription start sites lie within ±10 kb of transposable element–overlapping differentially accessible regions (TE-DARs; rhEPO vs. placebo). Enrichment results are shown for Biological Process and Cellular Component. Dot size indicates the number of genes associated with each term, and dot color represents statistical significance (−log₁₀ q-value). The enriched categories highlight pathways related to neurogenesis, synapse organization, neuronal signaling, ion transport, and membrane specialization, etc., reflecting functional processes linked to TE-derived regulatory remodeling under rhEPO treatment.

## SUPPLEMENTARY INFORMATION

**Supplementary Table S1**: Summary of sequencing and alignment metrics for snATAC–seq samples.

**Supplementary Table S2**: Transcription factor motif enrichment across Seurat-defined hippocampal clusters without treatment separation.

**Supplementary Table S3**: Transcription factor motif enrichment across the 11 major hippocampal lineages without treatment separation.

**Supplementary Table S4**: Summary of significant accessible chromatin peaks across the 11 hippocampal lineages and detailed annotation of peaks not present in the Mouse Brain Atlas.

**Supplementary Table S5**: Transposable element family enrichment within all accessible chromatin regions.

**Supplementary Table S6**: Globally defined differentially accessible regions, motif enrichments, TE-associated regulatory elements, and matched snRNA-seq comparisons for rhEPO and placebo conditions.

**Supplementary Table S7**: Correlation between differentially accessible regions and differentially expressed genes after filtering for significant values.

**Supplementary Table S8**: Significantly differentially accessible regions between rhEPO- and placebo-treated samples across the 11 hippocampal lineages.

**Supplementary Table S9**: Significantly differentially accessible regions between EPO-and placebo-treated samples across the 11 hippocampal lineages, annotated for proximal genes and overlapping transposable elements.

**Supplementary Table S10**: Gene regulatory network modules inferred for each of the 11 hippocampal lineages under EPO and placebo conditions.

**Supplementary Table S11**: Enrichment of transposable element families within differentially accessible regions identified between EPO- and placebo-treated samples.

**Supplementary Table S12**: Significant accessible chromatin peaks distinguishing newly formed from mature pyramidal neuron lineages.

**Supplementary Table S13**: Differentially accessible regions between EPO- and placebo-treated samples within newly formed and mature pyramidal neurons, annotated for nearby genes and transposable elements.

**Supplementary Table S14**: Enrichment of transposable element families within ChIP–seq peak sets for neurodevelopmental transcription factors.

**Supplementary Table S15**: Transposable element–associated differentially accessible regions located within 10 kb of annotated transcription start sites.

**Supplementary Table S16**: Gene Ontology analysis of genes associated with TE-overlapping differentially accessible regions bound by transcription factors.

**Supplementary Table S17**: List of neurogenic transcription factors included in TF-binding and enrichment analyses.

